# Ecological dynamics explain modular denitrification in the ocean

**DOI:** 10.1101/2024.09.25.615058

**Authors:** Xin Sun, Pearse Buchanan, Irene H. Zhang, Magdalena San Roman, Andrew R. Babbin, Emily Zakem

**Affiliations:** Department of Global Ecology, Carnegie Institution for Science, Stanford, CA, USA; Environment, Commonwealth scientific and industrial research organization, Hobart, Australia; Department of Earth, Atmospheric and Planetary Sciences, MIT, Cambridge, MA, USA; Department of Biological Sciences, University of Southern California, Los Angeles, CA, USA; Instituto de Biología Funcional y Genómica, IBFG-CSIC, Universidad de Salamanca, Salamanca, Spain

## Abstract

Microorganisms in marine oxygen minimum zones (OMZs) drive globally impactful biogeochemical processes. One such process is the multi-step denitrification, the dominant pathway for bioavailable nitrogen (N) loss and nitrous oxide (N_2_O) production. Denitrification-derived N loss is typically measured and modeled as a single step, but observations reveal that most denitrifiers in OMZs contain only subsets (“modules”) of the complete pathway. Here, we identify the ecological mechanisms sustaining diverse denitrifiers, explain the observed prevalence of certain modules, and examine the implications for N loss. We describe microbial functional types carrying out diverse denitrification modules by their underlying redox chemistry, constraining their traits with thermodynamics and pathway length penalties, in an idealized OMZ ecosystem model. Biomass yields of single-step modules increase along the denitrification pathway when growth is limited by organic matter (OM), explaining the viability of populations respiring nitrite and N_2_O in a nitrate-filled ocean. Results predict denitrifier community succession along environmental gradients: shorter versus longer modules are favored when OM versus N limits growth, respectively, suggesting a niche for the NO_3_^−^⟶NO_2_^−^ module in free-living communities and for the complete pathway in organic particles, consistent with observations. The model captures and mechanistically explains the observed dominance and higher oxygen tolerance of the NO_3_^−^⟶NO_2_^−^ module. Results also capture observations that nitrate is the dominant source of N_2_O. These results advance the mechanistic understanding of the relationship between microbial ecology and N loss, which is essential for accurately predicting the ocean’s future.

## Introduction

The ocean is experiencing rapid changes, including but not limited to warming (1) and deoxygenation (2). How marine microbes respond to changes in their environment feeds back on the biogeochemical cycles that control the availability of nutrients and the production of greenhouse gases, and thus influences the evolution of the oceanic ecosystem and Earth’s climate. Microbes that inhabit the anoxic layers of marine oxygen minimum zones (OMZs), located between the oxygenated surface and deep interior (3), have outsized biogeochemical impacts. Despite occupying less than 1% of the ocean’s volume, microbes in the anoxic zones of OMZs contribute up to 30% of the fixed nitrogen (N) loss from the ocean (4) to generate the widespread N deficit that limits primary production in over half of the surface ocean (5). As deoxygenation progresses (6), it is speculated that losses of fixed N may intensify and further constrain primary production at the surface (7), though dynamics remain uncertain (8). Greater N loss than N input from fixation may not only reduce the oceanic drawdown of atmospheric carbon dioxide but also increase the production of the potent greenhouse gas nitrous oxide (N_2_O), which is a key intermediate of anaerobic metabolism in OMZs (7, 9). Thus, understanding how diverse microbes interact and control N cycling within OMZs, especially N loss, is imperative for estimating the future of ocean health and climate.

Fixed N loss occurs through two microbial processes: anaerobic ammonium oxidation (anammox) and denitrification. Anammox (NH_4+_+NO_2_^−^⟶N_2_) is performed by a taxonomically conservative group of bacteria that fix carbon dioxide into their biomass using the energy generated from the anammox reaction. In contrast, denitrification is performed by diverse taxonomic groups. In the open ocean, denitrifiers are mostly heterotrophs that respire organic matter (OM) (10). Denitrification consists of multiple steps that sequentially reduce nitrate (NO_3_^−^) to produce reduced N (NO_3_^−^⟶NO_2_^−^⟶NO⟶N_2_O⟶N_2_). In practice, denitrification in OMZs is often assumed as a two-step reaction (i.e., NO_3_^−^⟶NO_2_^−^ and NO_2_^−^⟶N_2_), with nitrite (NO_2_^−^) as the intermediate excreted to and taken up from the extracellular environment, and thus N loss from denitrification is measured only as a single step (i.e., NO_2_^−^⟶N_2_) (11, 12). This one-step representation is incorporated into ocean biogeochemical models. However, recent metagenomic studies suggest that populations in OMZs have all possible combinations of genes from the denitrification pathway and that populations with only subsets of the full pathway are much more abundant than those containing the complete pathway (13–15). The mechanisms allowing for the prevalence of certain denitrification modules over others and the coexistence of modules competing for the same substrates remain unclear.

Because only some denitrifying modules result in N loss (Fig. 1a), intermediates may be rerouted towards or away from N loss pathways as the environment changes. Thus, microbial control of N loss is too complex to be accurately captured by single-step representation in ocean biogeochemical models. Trait-based ecosystem models resolving microbial functional types have the potential to synthesize and ultimately predict the outcome of these complex interactions. However, few denitrifiers in OMZs have been cultured or characterized, so we lack comprehensive knowledge of the quantitative traits and tradeoffs of the diverse metabolic strategies associated with different denitrifiers. Previous work has demonstrated that the chemical redox reactions underlying metabolism can be used to quantify traits of “metabolic functional types”, linking modeled microbial growth to the chemical potential of the environment in ways that may be more robust than relying solely on species- or location-specific measurements (16). This approach successfully captures coarse-grained biogeochemical dynamics in OMZs (17) and therefore has the potential to provide insight into the more complex dynamics of modular denitrification.

**Figure 1.**
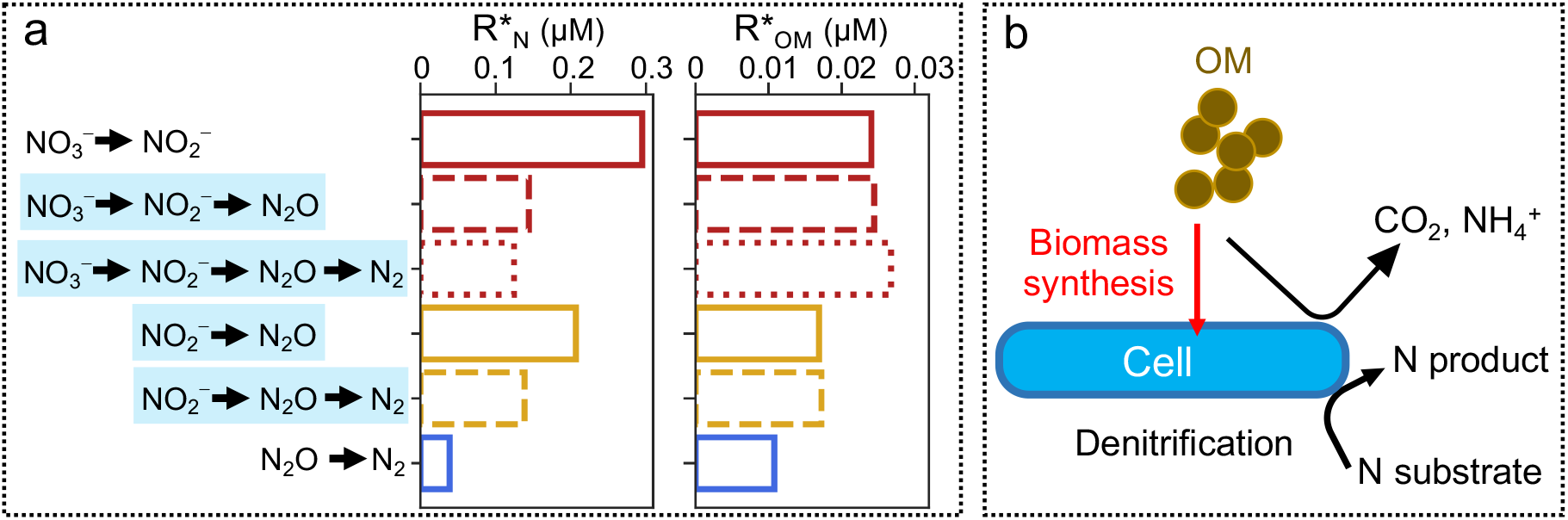
Schematic of denitrifier functional types. (**a**) The six modules of the denitrification pathway that are represented by microbial functional types in the ecosystem model, and their subsistence concentrations of OM (R*_OM_) and inorganic N (R*_N_) which reflect underlying thermodynamics of redox chemistry and proteome constraints via the biomass yields. A lower subsistence concentration allows microbes to be more competitive when the substrate is limiting (18) (see Methods). The colors of bars represent different N substrates, and line types of bars represent the number of denitrification steps of each functional type. (**b**) Schematic of the redox-fueled metabolism for a denitrifier cell. OM participates in both the biomass synthesis (anabolic) and the denitrification (catabolic) reaction. The balance of the energy needed by the former and generated by the latter sets the biomass yields.

In this study, we aim to understand how populations with diverse denitrification modules (hereafter all referred to as “denitrifiers” for simplicity) can co-occur in OMZs. As heterotrophs, all denitrifiers compete for OM, and some compete for the same N substrate, thus the observed diversity seems to contradict the competitive exclusion principle that the number of sustainable populations cannot exceed the number of resources (19, 20). We aim to identify the niches of different denitrifiers, exploring in particular why NO_3_^−^ ⟶NO_2_^−^ denitrifiers (often referred to as nitrate reducers) are the most abundant denitrifiers while complete denitrifiers are rare but still present. To answer these questions, we develop a new theoretical model that quantitatively describes functional types carrying out diverse denitrification modules by linking the redox reaction fueling each metabolism to traits (16) while accounting for proteome constraints for longer pathways (21–23) (Fig. 1).

We first examine the general patterns of how OM-vs. N-limited (i.e., electron donor vs. acceptor) growth regulates the modularity that emerges from thermodynamic constraints. We then incorporate the denitrifier functional types into an idealized OMZ ecosystem model, allow the ecosystem to self-assemble by resolving the interactions among functional types and substrates, and use resource-competition theory (18) to interpret the more complex patterns resulting from the interdependencies of the many types. We study the resulting succession of modularity along gradients in OM:NO_3_^−^ supply and oxygen supply. We show that diverse co-existing denitrifying microbes can be maintained by temporal and spatial niche partitioning and implicate this diversity as central to resolving key chemical features observed in OMZs.

## Results and discussion

### A theoretical framework for modular denitrification

We develop a theoretical framework to quantitatively describe microbial functional types carrying out different denitrification modules (Fig. 1a). We simplify the denitrification pathway as three steps (NO_3_^−^⟶NO_2_^−^⟶N_2_O⟶N_2_), neglecting the intermediate NO following previous work (24) because the genes producing and consuming NO are usually coupled (15, 25) and NO is unstable and the ability to consume it is a nearly-ubiquitous trait of microorganisms, beyond just denitrifiers (26). The three-step pathway allows for seven possible modules, including the six modules listed in Fig. 1a. The seventh, non-adjacent module (NO_3_^−^⟶NO_2_^−^ and N_2_O⟶N_2_) does not result in a potential niche within our framework, which suggests that an advantage for this type of metabolism remains unaccounted for, although its biogeochemical impact is represented by the corresponding single-step functional types (see Supplement). In general, a functional type may represent multiple species (or species-like equivalents) with the same function, and conversely, a metabolically flexible species may be associated with multiple functional types at different times.

We use the redox reaction fueling each module to estimate a metabolic budget for each functional type, resulting in a biomass yield for each required substrate and the stoichiometry for each excreted product. This metabolic budget thus links the biological (biomass of microbes) and chemical (nutrient concentrations) components in the model. Each functional type involves two coupled biomass yields: 1) the yield with respect to OM (*y*_*OM*_; mol biomass (mol OM)^−1^), which serves as the electron donor for the redox reaction and also provides building blocks for biomass synthesis, and 2) the biomass yield with respect to the reduced inorganic N species used as the electron acceptor (*y*_*N*_; mol biomass (mol N)^−1^). These biomass yields are coupled according to the fraction of electrons channeled towards biomass synthesis (anabolism reactions) vs. respiration (catabolism reactions), which is informed by the chemical potential (i.e., Gibbs free energy) of the reactions (27). We also apply pathway length penalties to the yields so that functional types carrying out more than one denitrification step are penalized to account for increased energetic costs associated with synthesizing more enzymes and transporters (28). Though the magnitude of the penalty is uncertain, we analyzed the qualitatively distinct outcomes of different penalty assumptions, choosing the assumption that results in a plausible outcome. The remaining uncertainty affects the model solutions quantitatively but not qualitatively (see Methods). To summarize, the resulting biomass yields of the denitrifier functional types differ because of two factors: the redox reaction - specifically, the stoichiometry and Gibbs free energy of the half-reaction for N reduction - and the denitrification pathway length. Our results are consistent with measured *y*_*N*_ for different N substrates of one cultured denitrifier (29), measured *y*_*N*_ for NO_3_^−^ of 75 taxonomically diverse denitrifiers with different denitrification steps (30) (Fig. S1b,d), and the estimated *y*_*OM*_ of a marine denitrifier (Marinobacter) in 59 different media by a new metabolic model (Fig. S1a,c).

The metric of competitive ability for functional types in the ecosystem is the resource subsistence concentration (18). In our model, the subsistence concentrations (R*_OM_ and R*_N_, Fig. 1a) are the minimum OM or N concentrations for which a functional type population is viable (i.e., when growth balances mortality) (18). Therefore, populations with a smaller R* can competitively exclude microbes with a larger R* when growth is limited by that resource and changes in its supply are slow relative to the timescales of microbial growth (16, 18). Here, R*_OM_ and R*_N_ of denitrifiers reflect solely the biomass yields (Equation 3 in Methods) with higher yields corresponding to lower R*. Though differences between R* values are small, our premise is that their ordering is robust following the fundamental constraints. Thus, we introduce no other distinctions between the functional types: we parameterize each denitrifier type with the same biomass loss rate (i.e., the dilution rate of the virtual chemostat) and the same uptake kinetics for the same substrate. In other words, we assume that the underlying constraints to the functional types rather than their ability to evolve their substrate affinity and other traits, in a given environment, predominantly differentiate their fitness.

### General patterns of competitive ability for OM and N resulting from the theoretical framework

The resulting biomass yields order the competitive abilities of the denitrifier functional types for OM and N (Fig. 1a & Fig. S1a,b). For the three single-step modules (pathway length = 1), the efficiency of both OM and N use increases along the pathway, so that R* for both OM and N are highest (least competitive) for NO_3_^−^⟶NO_2_^−^, lower for NO_2_^−^⟶N_2_O, and lowest (most competitive) for N_2_O⟶N_2_. This means that if these three functional types were all competing for OM alone (i.e., if NO_3_^−^, NO_2_^−^, and N_2_O were all abundant), the N_2_O⟶N_2_ type would outcompete the other two. However, as explored in the ecosystem model below, subsequent modules may often rely on earlier modules to supply the N.

For longer pathways, when considering only reaction thermodynamics (without the pathway length penalty), the efficiency of both OM and N use increases with pathway length, since more energy is extracted from the same electron donor. Therefore, complete denitrifiers would always competitively exclude all other types. However, this is in stark contrast to observations that complete denitrifiers are rare within OMZs. If we consider the cost of maintaining a longer pathway, the efficiency of OM use instead decreases for longer pathways, while the efficiency of N use still increases (Fig. 1a & Fig. S1a,b). Thus, the complete denitrification module is least competitive for OM, but most competitive for NO_3_^−^. This is a robust result across the plausible penalty space used in our framework (Methods) and is consistent with estimated yields using a metabolic model in 59 different media (Fig. S1a,c) and measured yields in 75 taxonomically diverse denitrifiers (30) (Fig. S1b,d).

Together, the constraints from thermodynamics and pathway length penalty suggest a set of general patterns in competitive ability: 1) when functional types are limited by OM, shorter denitrification pathways can competitively exclude longer pathways using the same N substrate, and 2) when functional types are limited by the same N substrate, longer denitrification pathways can competitively exclude shorter pathways. These patterns reflect a tradeoff between pathway length and yield, and in this way are similar to the well-studied cellular optimization for pathway length when considering only organic carbon limitation: the longer pathway (respiration) has a higher yield, and so is favored over the shorter pathway (fermentation) when organic carbon is limiting (21–23, 31). These patterns are also consistent with the hypothesis that complete ammonia oxidizers (comammox) performing two steps of nitrification in one cell is favorable, compared to the division of labor between two microbial groups in an N-limited environment, again because the longer pathway has a higher yield but is penalized by pathway length (28, 32, 33). Here, we explore the more complex patterns emerging when considering requirements for two different resources (OM and N).

These patterns provide guiding hypotheses for how the denitrification modularity is ordered by substrate availability. We hypothesize the prevalence of the NO_3_^−^⟶NO_2_^−^ module (the observed dominant module) in the NO_3_^−^-rich, OM-limited open-ocean OMZ environment (10, 11, 34) over complete denitrification. We next use these patterns and hypotheses from the theoretical analysis to understand more complex outcomes from the interactions of the functional types in the ecosystem model where some modules depend on others for required intermediates.

### The coexistence of single-step modules

The substrate of the first denitrification step (NO_3_^−^⟶NO_2_^−^), NO_3_^−^, is the most abundant form of fixed N in the ocean. Consequently, the latter steps of denitrification usually rely on prior steps to produce their N substrates. Because NO_3_^−^ is rarely consumed to completion in open-ocean marine OMZs, one might ask why microbes would evolve to develop the cellular machinery to reduce less available N resources such as NO_2_^−^. The fact that microbes have evolved this capability is well established, so this is a theoretical question. Our model frames the underlying energetics in terms of resource competition (i.e., R*, Fig. 1a), and thus provides a new mechanistic explanation for the use of N intermediates from an ecological perspective.

We incorporate the three functional types carrying out the single-step modules into a simple ecosystem model. We supply OM and NO_3_^−^ at a ratio that results in OM-limited growth. Initially, the NO_3_^−^ reducer is the only functional type that can be sustained. Its production of NO_2_^−^ then allows the NO_2_^−^ reducer to invade, becoming ecologically sustainable because of the NO_2_^−^ reducer’s superior competitive ability for OM (i.e., lower R*_OM_). However, the NO_2_^−^ reducer’s reliance on the NO_3_^−^ reducer for NO_2_^−^ means that it becomes NO_2_^−^-limited, allowing the NO_3_^−^ reducer to continue to survive. Similarly, the N_2_O reducer is then able to invade because it is the superior competitor for OM, but it also then becomes N_2_O-limited and so cannot competitively exclude the other types. Therefore, all three types carrying out the single-step modules coexist at steady states (Fig. S2a) due to the dependence of the latter modules on the former.

Thus, our analysis combines the thermodynamics of denitrification with ecological theory to provide a fundamental explanation for why populations have evolved to carry out modules using N species other than NO_3_^−^ as oxidants. The ecological model clarifies that populations carrying out denitrification modules using intermediate substrates (i.e., NO_2_^−^ and N_2_O) are viable when OM is limiting only because they are sequentially better competitors for OM. If the thermodynamics were such that the latter modules along the pathway were associated with a lower generation of free energy, the latter modules would not be able to invade the ecosystem and sustain their populations.

### Community succession along the OM:NO_3_^−^ supply gradient

We next introduce more functional types into the ecosystem model while varying the ratio of OM to NO_3_^−^ supply to simulate different ocean environments. This OM:NO_3_^−^ supply spans the OM-limited, open-ocean conditions encountered by “free-living” microbes to the OM-rich conditions inside or nearby organic particles (35). We solve for the steady state at each supply ratio. Though the results are complex, they remain interpretable using our two general patterns for OM vs N limitation and pathway length: OM competition favors shorter pathways, and N competition favors longer pathways.

First, we illustrate these patterns in the simplest way possible by considering only the three functional types that use NO_3_^−^ (Fig. 2a,d,g). As OM supply rate increases relative to NO_3_^−^ supply, the system shifts from OM limitation to NO_3_^−^ limitation (Fig. 2g), and the pathway length of the surviving denitrifier type increases (Fig. 2d). Thus, as hypothesized by the general patterns, single-step NO_3_^−^⟶NO_2_^−^ is dominant when OM is limiting, while complete denitrification is dominant when NO_3_^−^ is limiting, and two-step(NO_3_^−^ ⟶NO_2_^−^⟶N_2_O is sustained at intermediate supply (Fig. 2a). The thresholds that delineate the prevalence of each functional type and the stable coexistences among them can be computed from the traits (yields) using resource competition theory (36). Specifically, the three critical OM:NO_3_^−^ supply ratios separating the four distinct survival regimes are defined by the reciprocal of the consumption vectors of the three functional types (Fig. S3), which reflect *y*_*NO3*_:*y*_*OM*_ (grey lines in Fig. 2a).

**Figure 2.**
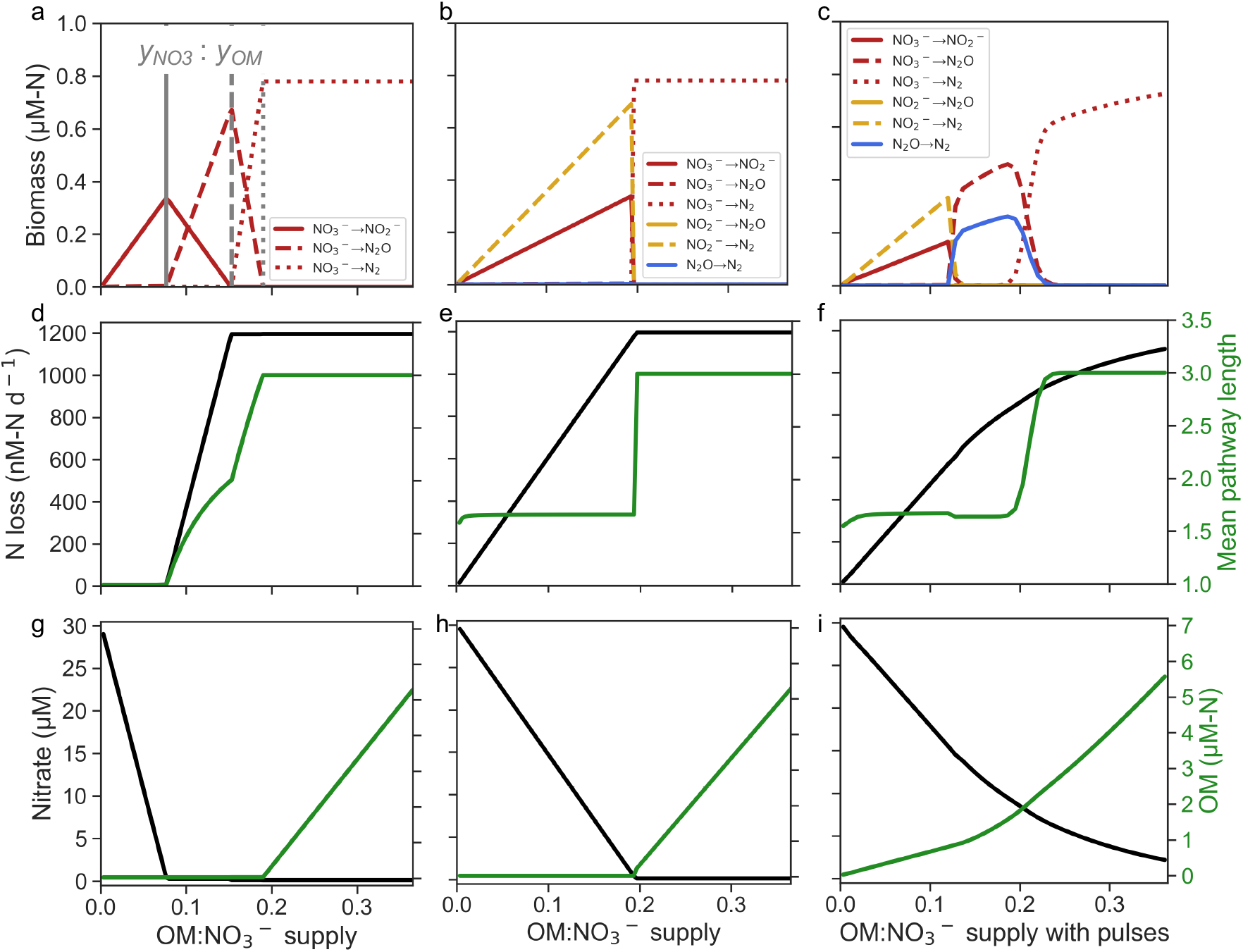
The shift of denitrifier community composition (**a, b, c**), N loss via denitrification and mean denitrification pathway length (**d, e, f**), and equilibrium (with constant OM supply) or quasi-equilibrium (with OM pulses) NO_3_^−^ and OM concentrations (**g, h, i**) with increasing OM:NO_3_^−^ supply (mole-N (mole-N)^−1^) when the O_2_ supply rate is 0. Results from the model including (**a, d, g**) three NO_3_^−^-reducing denitrifier functional types with constant OM supply, six denitrifier types with (**b, e, h**) constant OM supply, and (**c, f, i**) OM pulses. Vertical grey lines in (**a**) are reciprocal of consumption vectors (1/CV = *y*_*NO3*_/*y*_*OM*_) of the three denitrifiers (Fig. S3). The N loss in black (**d, e, f**) is the reduction rate of any fixed N to any gaseous N as shown in Fig. 1a. The mean pathway length in green (**d, e, f**) is the sum of the pathway length of each denitrifier population weighted by their biomasses.

Next, we include six denitrifier functional types in the model and again solve for the steady state along the OM:NO_3_^−^ supply gradient (Fig. 2b,e,h). While the increase in OM supply rate shifts the nutrient regime generally from OM limitation to NO_3_^−^ limitation (Fig. 2h), the cross-feeders using NO_2_^−^ and N_2_O are N-limited for the entirety of the domain. This complex network of cooperation and competition results in a community succession along the OM gradient. When OM supply is lowest, single-step NO_3_^−^⟶NO_2_^−^ and two-step NO_2_^−^⟶N_2_O⟶N_2_ are sustained (Fig. 2b & Fig. S2b). This is because OM limits the types competing for NO_3_^−^, favoring the shorter pathway, but the NO_2_^−^ reducers are limited by NO_2_^−^, favoring the longer pathway beginning with NO_2_^−^. In the OM-rich, NO_3_^−^-limited regime, the three-step complete denitrifier is sustained. This suggests that OM-rich environments such as particles (35, 37) or shallower depths with high OM supply (38) are potential niches for the rare and otherwise energetically unfavorable complete denitrifiers.

Note that the model distinguishes the ratios of biomasses from the ratios of associated N transformation fluxes according to the redox stoichiometry (Fig. S4). For example, at low OM:NO_3_^−^, the biomass of the NO_3_^−^⟶NO_2_^−^ type is lower than that of the NO_2_^−^⟶N_2_O⟶N_2_ type, but the NO_3_^−^⟶NO_2_^−^ transformation rate is equal to the NO_2_^−^⟶N_2_ transformation rate. This exemplifies that ratios of gene abundances, which relate more directly to biomass ratios, may be different than ratios of measured rates.

### Time-varying substrate supply creates an intermediate regime

Analyses thus far have assumed a steady supply of OM and NO_3_^−^ at each point along the supply ratio gradient. The resulting steady state enables complete competitive exclusion, according to R*_OM_ and R*_N_. However, phytoplankton blooms, eddies, extreme weather, and subduction events mean that substrate supply, particularly OM supply, varies in time (39–42). Moreover, the reworking of heterogenous OM by different heterotrophic communities changes the availability of particular organic substrates over time (34). To mimic this temporal variability and examine how it impacts coexistence regimes of denitrifiers, we supply OM in pulses to the virtual chemostat as instantaneous OM additions at a certain frequency (Methods).

Time-varying OM supply results in a more gradual transition from OM-limited to NO_3_^−^-limited state (Fig. 2i & Fig. S4k). Critically, time-varying supply introduces an intermediate regime (with two-step NO_3_^−^ ⟶NO_2_^−^⟶N_2_O and single-step N_2_O⟶N_2_ types) where N_2_O is the intermediate substrate (Fig. 2c & Fig. S4). Observations show that the former type (NO_3_^−^⟶NO_2_^−^⟶N_2_O) is the major contributor to N_2_O production in OMZs (43–45). Our model suggests that non-steady state conditions represented by time-varying OM supply here allow the prevalence of this functional type (Fig. 2b,c).

The frequency of OM pulses affects the coexistence pattern of denitrifiers quantitatively but not qualitatively (Fig. S4c,d), with higher frequencies approximated by a constant supply. The relevant timescales in open-ocean OMZs are that of the environmental perturbations relative to that of microbial growth. OM pulses associated with large-scale physical processes in the open ocean usually occur less frequently (weekly to monthly) (46) than microbial turnover (every few hours to days) (47). This timescale ratio is consistent with our choice of OM frequency relative to the dilution rate of the chemostat model in Fig. 2, suggesting that our intermediate regime characterized by the NO_3_^−^⟶NO_2_^−^⟶N_2_O type may indeed be common in OMZs.

### Higher oxygen tolerances of NO_3_^−^ reducers with cryptic oxygen cycling

So far, analysis has exclusively considered the anaerobic denitrification metabolisms. However, denitrifiers in OMZs occupy both functionally anoxic layers and oxic-anoxic interfaces, and it is also becoming increasingly clear that “cryptic” oxygen cycling is a key feature of the seemingly anoxic layers where small but non-negligible amounts of oxygen are produced locally or delivered through physical processes (48–52). Therefore, denitrifying populations subsist in environments where trace oxygen is supplied, even though oxygen may not accumulate. Thus, the interactions of anaerobic and aerobic functional types may impact community structure and denitrification modularity.

To examine this impact, we add an axis of oxygen supply to the model (Fig. 3). We include three key aerobic functional types: aerobic heterotrophs, ammonia-oxidizing archaea (AOA), and NO_2_^−^-oxidizing bacteria (NOB) using recently published parameterizations that are consistent with observations (53, 54). Following observations, we assume that chemoautotrophic functional types have a higher affinity for inorganic N than the denitrifiers, who must devote a portion of their cellular machinery to OM uptake and processing (55–58). We also add an anaerobic ammonia-oxidizing (anammox) functional type to complete the set of functional types required to represent key N cycling and loss processes in open-ocean OMZs. As in previous work (17), we do not impose any oxygen thresholds or oxygen tolerances on microbial functional types and rather examine the relationships with oxygen that emerge from the ecological interactions. Consistent with the previous work (17), anaerobic metabolisms are sustained at a sufficiently low ratio of oxygen to OM supply (i.e., *ϕ* = 1, the white line in Fig. 3a).

**Figure 3.**
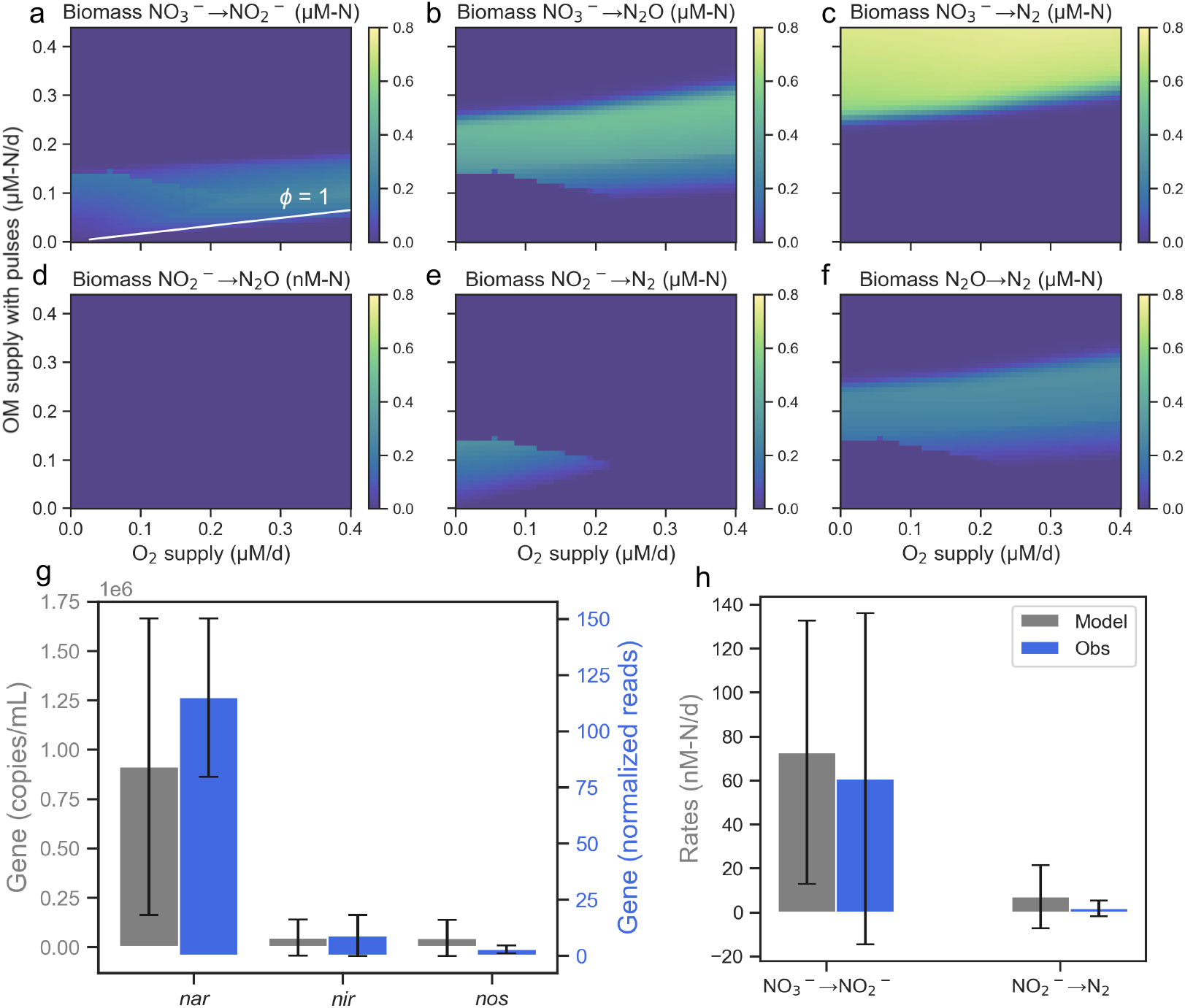
Denitrifier biomasses estimated by the model with OM pulses (**a-f**) and the comparison between modeled and observed gene abundance and rates (**g, h**). The white line in **a** indicates where *ϕ* = 1. (**g, h)** Modeled gene abundances of *nar* (nitrate reductase genes encoding NO_3_^−^⟶NO_2_^−^), *nir* (nitrite reductase genes encoding NO_2_^−^⟶NO) and *nos* (nitrous oxide reductase genes encoding N_2_O⟶N_2_), and the modeled rates of NO_3_^−^⟶NO_2_^−^ and NO_2_^−^⟶N_2_ when a mean OM supply rate is 0.015 µM-N/d and when quasi-equilibrium O_2_ concentration is below 2 µM (the O_2_ detection limit of the Seabird sensor used to measure the rates in the figure) and measured gene abundances (13) and rates (59) in the Eastern Tropical North Pacific OMZ when O_2_ concentration is below the Seabird sensor detection limit (2 µM). The model assumes cells have four copies of *na*r (13, 60) and one copy of *nir* or *nor*. Both modeled and observed gene abundances reflect relative abundances of genes, and thus the order instead of the absolute number of the genes are comparable. Error bars are standard deviations.

Results show that modeled denitrification modularity varies with oxygen supply, despite assuming no direct oxygen sensitivity of denitrification (Fig. 3a-f). Specifically, results suggest that NO_3_^−^-reducing denitrifiers are able to survive under a wider range of oxygen conditions than NO_2_^−^-reducing denitrifiers (Fig. 3a,b,c,e, Fig. S5 & S6), matching observations (43, 59, 61, 62). The apparently stronger oxygen inhibition on NO_2_^−^-reducing denitrifiers relative to NO_3_^−^-reducing denitrifiers in our model is because the former become limited by NO_2_^−^ when oxygen is available due to the presence of aerobic NOB. NOB outcompete the NO_2_^−^-reducing denitrifiers due to their higher affinity for NO_2_^−^ (56, 58, 63). NOB’s high affinity for oxygen also allows them to grow at low oxygen supply (64, 65). This outcome is for the case when NO_2_^−^ is limiting but oxygen supply is non-negligible, such as in the low oxygen waters surrounding an anoxic core. The complex interactions produce a nonmonotonic decrease in the NO_2_^−^⟶N_2_O⟶N_2_ functional type with increasing oxygen (Fig. 3e). A model sensitivity experiment without the NOB functional type shows that this nonmonotonic decrease disappears when the complex interaction between denitrifiers and NOB is excluded (Fig. S7), and additional model sensitivity tests demonstrate the robustness of this mechanism to the most uncertain parameters (Supplement). This result is also intuitive: the apparent higher oxygen tolerance of NO_3_^−^ reducers emerges because NO_3_^−^, unlike NO_2_^−^, is always available when oxygen is present. These results hypothesize fundamental explanations from the perspective of microbial niche differentiation for why some OMZ microbes have not evolved away from actual, measured oxygen sensitivities that reflect oxygen-intolerant enzymes.

### Dominance of the NO_3_^−^⟶NO_2_^−^ module

Synthesizing our analysis of variations in both OM:NO_3_^−^ and oxygen supply in the ecosystem model provides an explanation for why NO_3_^−^⟶NO_2_^−^ is the most abundant denitrifying pathway observed in OMZs (13–15). First, most of the ocean is considered oligotrophic in terms of OM supply (10), resulting in the modeled predominance of NO_3_^−^⟶NO_2_^−^ and NO_2_^−^⟶N_2_ modules when OM supply is low (Fig. 2). Second, when adding the wider oxygen niches of types using NO_3_^−^ (Fig. 3 & Fig. S6), the prevalence of the NO_3_^−^ ⟶NO_2_^−^ module over NO_2_^−^⟶N_2_ is predicted. We convert modeled biomasses to a quantity that is proportional to gene abundance by summing up the biomass of all microbial functional types that would contain a certain gene and assuming one gene copy per cell with the exception of *nar* based on observations (13, 60). These estimated gene abundance proxies are consistent with measured gene abundances showing that the gene encoding the NO_3_^−^⟶NO_2_^−^ pathway (*nar*) is significantly more abundant than the gene encoding NO_2_^−^⟶NO (*nir*) (Fig. 3g). Rates of NO_3_^−^⟶NO_2_^−^ are much higher than the rates of NO_2_^−^⟶N_2_ in both model and data (59, 61, 66) (Fig. 3h).

### Controls on N loss

The modeled denitrifier community succession suggests accompanying patterns in N cycling and loss dynamics. As the dominant denitrifiers shift with OM:NO_3_^−^ supply, there is also a shift in the dominant N species predicted by the model to be produced, excreted from the cell, and then consumed by different cells (Fig. 4), which we here term the N “intermediates”. These intermediates may not accumulate, because they may be consumed as quickly as they are produced. Thus, an N species’ status as an intermediate in our framework is a necessary but not sufficient condition for its accumulation. At low OM:NO_3_^−^ supply, NO_2_^−^ is the only N intermediate. This result is consistent with the general understanding that NO_2_^−^ is a key intermediate in OM-limited open-ocean OMZs. At relatively higher OM:NO_3_^−^ supply, the regime that emerges from time-varying dynamics, N_2_O is the intermediate. This result suggests that active N_2_O fluxes, and thus the potential for denitrification-derived N_2_O emissions from ocean to atmosphere, require relatively higher OM:NO_3_^−^ supply and a more dynamic environment than many oligotrophic open-ocean locations. Because OM supply generally wanes with depth, this implies that this denitrifier-led N_2_O cycling may be most critical in the dynamic areas of the upper OMZ oxycline in productive locations. This is consistent with the most intense N_2_O accumulation and N_2_O cycling rates being measured in the upper oxycline (45, 67– 69). At the highest OM:NO_3_^−^, likely inside organic particles (35) where N-limitation favors complete denitrification. This suggests a smaller role for N_2_O cycling and for anammox (which requires NO_2_^−^) inside or in close association with particles.

**Figure 4.**
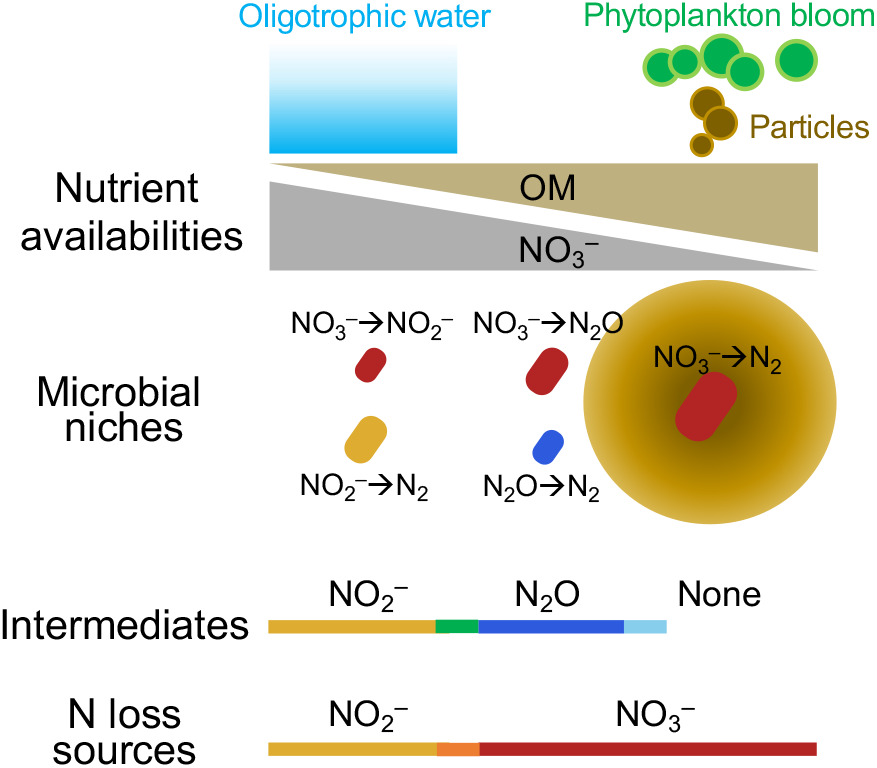
Schematic of microbial niches, intermediates of the denitrification process, and N loss sources under different nutrient conditions predicted from our theoretical framework and model. The green and light blue bars in the “intermediates” panel indicate niche overlaps of microbes resulting in NO_2_^−^ and N_2_O cycling, N_2_O and no intermediate, respectively. The orange bar in the “N loss sources” panel indicates a niche overlap of microbes resulting in NO_2_^−^ and NO_3_^−^ being N loss sources.

The community transition also shifts the bioavailable N that is directly transformed (within one functional type) into a gaseous form (N_2_O or N_2_), which we term the “N loss source” (Fig. 4). At low OM:NO_3_^−^ supply, NO_2_^−^ is the main N loss source from denitrification. Since anammox requires NO_2_^−^, this regime is the only regime compatible with anammox being an active and dominant contributor to N loss. Indeed, in the model, the contribution of anammox to N loss is consistent with previous theory (∼30%) (11, 70, 71) when aerobic heterotrophy is negligible (since increasing oxygen supply means that more OM is respired aerobically, but ammonia supply from heterotrophs remains about the same). This implies that anammox is sustainable when denitrification is limited by OM rather than NO_3_^−^ (Fig. S5, black line).

In contrast, as OM:NO_3_^−^ supply increases, NO_3_^−^ rather than NO_2_^−^ becomes the N loss source. This suggests that measurements targeting NO_2_^−^ as the source of N loss in OM-rich environments may be missing much of the flux. In this way, results provide a set of testable hypotheses for considering N loss from the ocean for new experiments and observations.

Overall, the succession of denitrifier communities along environmental gradients results in a succession of active intermediates and N loss sources of denitrification. In general, in less productive waters where OM supply is lower, NO_2_^−^ cycling is more favorable. In contrast, N_2_O cycling by denitrifiers is predicted to be generally more favorable in more productive, anoxic locations where OM supply is dynamic. It is important to note that our results do not imply that NO_2_^−^ and N_2_O cycling must be spatially separated. They can occur in the same water mass due to inherent complexities allowing different subsets of denitrifiers to experience different OM:NO_3_^−^ supply. For example, free-living (low OM:NO_3_^−^ supply) and particle-associated (high OM:NO_3_^−^ supply) communities may be sampled together. Additionally, different denitrifier species may preferentially consume different organic substrates, and the relative supply of OM:NO_3_^−^ differs for these diverse types of OM. Our simple model provides the necessary first step of untangling this complexity and can aid in the interpretation of more complex regimes. The broad, simple patterns presented here can provide guiding principles and testable predictions that can be used to design sampling strategies targeting different fractions of the same water mass or environmental gradients to better understand the relationship between complex microbial ecology and N cycling.

### Concluding remarks

Models simplify complex natural ecosystems based on what questions they are designed to answer. Many biogeochemical models without microbial components have captured key biogeochemical features of OMZs (24, 72–74). However, our newly developed theoretical model with explicit microbial functional types is necessary to explain the observed functional diversity of denitrifying microbial populations and to elucidate the biogeochemical dynamics arising from their interactions with each other and with the rest of the ecosystem.

Our framework suggests that the coexistence of microbes in OMZs can largely be attributed to temporal and spatial niche partitioning in accordance with ecological theories and experiments (20, 36, 75, 76). This framework reproduces key observed features of the N cycle in OMZs, including the dominance of the NO_3_^−^⟶NO_2_^−^ pathway, particularly in oligotrophic waters more prone to oxygen supply, and its higher oxygen tolerance. Additionally, it captures the dominance of NO_3_^−^⟶NO_2_^−^⟶N_2_O pathway in N_2_O production. While microbial functional types driving different denitrification pathways may belong to distinct niches in our framework (Fig. 4), their coexistence is possible through micro-scale gradients in both space and time, which would allow diverse denitrification processes to occur and intermediates (NO_2_^−^ and N_2_O) to accumulate in what is observed to be the same water mass. Other environmental or biological factors, such as eddies and mixing, the top-down control of biomass by grazing or viral lysis, the variety of OM substrates as distinct resources, and additional tradeoffs among traits that characterize other fitness aspects of microbial functional types (75) can also influence the diversity of denitrifiers.

Our results also produce testable hypotheses for niche partitioning and its relationship with N loss dynamics. The model suggests that denitrifier functional types performing longer denitrification pathways thrive under N limitation conditions but may become outcompeted by types performing shorter pathways once OM becomes limiting. Consistent with gene-level (13, 77) or genome-resolved metagenomic observations (15), our results show that the most abundant functional type, the NO_3_^−^⟶NO_2_^−^ module, prefers a free-living lifestyle, and the rare functional type, complete denitrification, becomes competitive in OM-rich, N-limited particles (Fig. 4). This niche differentiation maps onto a hypothesized shift of denitrification intermediates and N loss sources when nutrient limitation changes (Fig. 4). Our framework, for example, indicates that relatively high levels of OM supply in a dynamic, low-oxygen environment will favor extracellular N_2_O fluxes from denitrification, which, if conditions allow, may accumulate and then potentially then fuel N_2_O emission to the atmosphere. These conditions may be more commonly satisfied in the upper OMZ, close to the oxycline, where inputs of OM are strong and variable. Also, results suggest that the source of N loss, from outside the cell, will switch from NO_2_^−^ to NO_3_^−^ with increasing OM supply (i.e., from free-living to particle-associated environments). These hypotheses can be tested using single-cell or metagenomic sequencing and stable isotope incubation experiments.

This simple framework, rooted in thermodynamics and redox chemistry, could be incorporated into higher-dimension models or extended to other ecosystems. With the explicit, yet theory-based representation of microbes, our model sets the stage for considering the effect of microbial eco-evolutionary dynamics on ocean biogeochemistry, which is critically needed to understand and predict the response of ecosystems to global change.

## Methods

### Denitrifier functional types

Our model framework quantitatively describes diverse denitrifier functional types in an ecosystem model of an anoxic zone based on their unique redox chemistry. This framework builds on previous redox-based frameworks for aerobic and chemoautotrophic functional types (17). We simplify the denitrification pathway into NO_3_^−^⟶NO_2_^−^⟶N_2_O⟶N_2_. All possible combinations of denitrification steps result in seven possible functional types (Fig. 1a): three NO_3_^−^-reducing denitrifier types (NO_3_^−^⟶NO_2_^−^, NO_3_^−^⟶NO_2_^−^⟶N_2_O, NO_3_^−^⟶NO_2_^−^⟶N_2_O⟶N_2_), two NO_2_^−^-reducing denitrifier types (NO_2_^−^⟶N_2_O, NO_2_^−^⟶N_2_O⟶N_2_), one N_2_O-reducing denitrifier types (N_2_O⟶N_2_), and one group with non-adjacent denitrification steps (NO_3_^−^⟶NO_2_^−^ and N_2_O⟶N_2_) (15). This latter non-adjacent type is competitively excluded in a model simulation (Supplement).

### Thermodynamics methodology

We estimate biomass yields *y*_*OM*_ and *y*_*N*_ from thermodynamics and pathway length penalty. First, we use redox half reactions to describe and summarize the partitioning of electrons to balance biomass synthesis (anabolism) with energy-generating redox reactions (catabolism) (See Supplement for details). These half reactions do not necessarily represent the detailed reactions occurring at the enzyme level. The standard molal Gibbs free energy of formation (Δ*G*^0^) values for compounds in the model as a function of temperature are from (78). We calculate the actual Gibbs free energy (Δ*G*) from (Δ*G*^0^) using the Nernst equation, considering the ballpark concentrations of substrates measured in OMZs. The total energy needed for synthesizing biomass per electron transferred (Δ*G*_*s*_) is defined as the energy needed to convert OM to pyruvate plus the energy needed to convert pyruvate to biomass considering an efficiency term of electron transfer (*ep*). The energy released per electron from the redox reaction (Δ*G*_*r*_) is different for each denitrification reaction. Following Rittmann & McCarty (27), we estimate the fraction *f* of electrons from OM used for biomass synthesis versus the fraction used in the energy generating redox reaction (i.e., denitrification). Assuming the same inefficiency *ep* for the energy production, the energy produced from denitrification (Δ*G*_*r*_ · *ep* · (1 − *f*)) must equal to the energy needed for biomass synthesis (−Δ*G*_*s*_ · *f*). Thus, 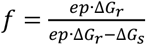 (Supplement).

Second, a penalty (P) is applied to denitrifiers possessing more than one denitrification step assuming that higher proteome costs associated with longer steps require more electrons from OM to be used in energy-generating redox reactions (Fig. 1b). Because the functional form and magnitude of such a penalty is not clear, we here assume a linear relationship between the penalty and the pathway length, but other functional forms would give the same results qualitatively. The penalty is applied to the fraction of electrons used for biomass synthesis: *f* · (1 − *P* · (*n* − 1)), where n is the number of denitrification steps (the pathway length). We analyze the effect of different P values on the *y*_*OM*_ and *y*_*N*_ of different denitrifiers (Supplement) because measurements or other estimates of *P* are limited (Fig. S8). The uncertainties in our penalty parameterization, as well as uncertainties in other functional type parameters (e.g. maximum uptake rate of OM, half-saturation constants, and loss rates), will impact the specific values of *P*_*C1*_ and *P*_*C2*_. However, the same scenarios presented here will dictate the ecological dynamics. Thus, our results are qualitatively robust to uncertainties and variations in parameters and penalty parameterizations.

### Our estimated yields supported by experimental observations and metabolic modeling

To validate our estimated biomass yields *y*_*OM*_ and *y*_*N*_ from thermodynamics and pathway length penalty, we compared them with independent observations from measurements and results from a metabolic model. The first comparison was with *y*_*N*_ values for different N substrates (specifically NO_3_^−^, NO_2_^−^, and N_2_O) in *Pseudomonas denitrificans* grown in a chemostat culture, where glutamate served as the source of energy, carbon, and N (29). The second comparison was with measured *y*_*N*_ values for NO_3_^−^ from 75 taxonomically diverse denitrifiers (including alpha, beta, and gamma *Proteobacteria*) under batch culture conditions with succinate as the carbon source (30). Our estimated *y*_*N*_ is consistent with these experimental measurements (Fig. S1b,d).

Additionally, to validate the *y*_*OM*_ values obtained from our theoretical framework, we compared them with estimates derived from metabolic modeling (Supplement). We reconstructed the metabolic model of a marine denitrifier (*Marinobacter*) and used Flux Balance Analysis (79) to estimate *y*_*OM*_ across 59 different media, each with a distinct carbon source. The results from metabolic modeling are also consistent with our *y*_*OM*_ estimates (Fig. S1a,c).

### Functional type growth

The growth rate (*µ*) of any given functional type on a required resource is set by the specific resource uptake rate and the biomass yield with respect to growth on that resource, following previous methods (17), as:

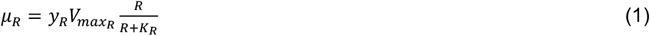

where the resource uptake rate (*V*_*R*_) for any resource *R* is modeled by the Michaelis-Menten equation (80). We assume identical uptake kinetics (*V*_*max*_ and *K*_*R*_) for each resource for all denitrifier functional types in order to isolate the impacts of thermodynamics and pathway length penalty. The modeled growth rate is defined by the limiting growth rate for multiple required resources (i.e., OM and N for the denitrifiers), following Liebig’s Law of the minimum, as:

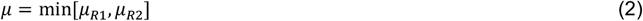

The subsistence concentration of a resource (*R\**) that is required for microbial growth is the concentration that makes the growth rate of the microbial population equal its loss rate (which is the dilution rate for the chemostat model; *µ = D*):

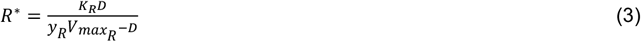

### Virtual chemostat model

The equations below describe changes in microbial biomass and nutrient concentrations over time in the virtual chemostat, following previous methods (17, 53). OM, NO_3_^−^, and oxygen are supplied to the chemostat according to prescribed incoming concentrations [OM]_in_, [NO_3_^−^]_in_, and [O_2_]_in_ (mmol m^−3^). *D* is the dilution rate of the chemostat (day^−1^), *y* is the functional type and substrate-specific yield (mol biomass N synthesized per mol substrate consumed), *e* is the excretion ratio (moles of a molecule excreted per mole biomass N synthesized, proportional to *y*^*-1*^). Both *y* and *e* are set by the stoichiometry of the redox reactions listed above for the denitrifiers, and as described in references (17, 54) or the additional aerobic and chemoautotrophic functional types. *µ*_*i*_ (day^−1^) is the growth rate of functional type *i*, and *B*_*i*_ (mmol N m^−3^) is the biomass. All inorganic N and OM concentrations are normalized to mmol N m^−3^. For example, the concentration of N in the form of N_2_O in the model (N_N_2_O) is twice that of N_2_O.

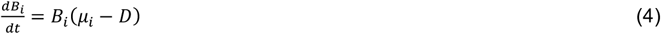

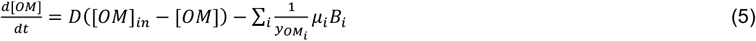

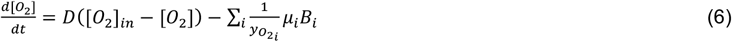

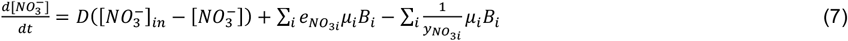

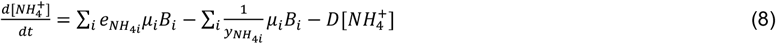

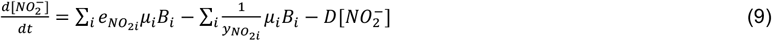

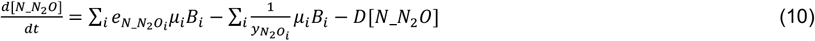

We solve Eqns. 4 - 10 for variable OM:NO_3_^−^ supply. We vary [OM]_*in*_ and assume a constant NO_3_^−^ supply of 30 µM to simulate the NO_3_^−^-filled ocean. The equations are solved numerically by integrating forward in time until a dynamic equilibrium is reached.

### Functional type survival and coexistence regimes

We use Resource Competition Theory (36) to quantitatively identify the survival and coexistence regimes for the three NO_3_^−^-reducing denitrifiers. These three types compete for two non-substitutable resources (NO_3_^−^ and OM). On this two-resource plane, we can draw a zero net growth isocline (ZNGI) for each type under NO_3_^−^ or OM limitation (Fig. S3), which is R*_N_ or R*_OM_, respectively. A consumption vector (CV), the ratio of the two resources being consumed, is calculated for each functional type as:

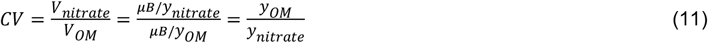

The CVs of the three NO_3_^−^-reducing denitrifiers indicate the critical OM:NO_3_^−^ supply ratios that set the survival and coexistence regimes.

## Acknowledgments

We would like to thank Dr. Clara Fuchsman and Dr. John Tracey for providing their valuable published observation data. We appreciate Dr. Daniel McCoy’s insightful comments on our manuscript. Xin Sun was supported by the Simons Foundation Postdoctoral Fellowship in Marine Microbial Ecology. Irene Zhang was supported in part by an MIT School of Science MathWorks Science Fellowship. ARB was supported by NSF grants OCE-2138890 and OCE-2142998. EJZ thanks NSF (Grant #2125142) and Carnegie Science for funding.

## Code availability

All code, model parameters and the reconstructed metabolic model are deposited in GitHub (https://github.com/xinsun12/ChemostatModel_ModularDenitrification_clean) and will be linked to a DOI using Zenodo when accepted for publication.

