## Supplementary materials for "Ecological dynamics explain modular denitrification in the ocean"

### The explicit representation of microbial functional types in a model framework

*Stoichiometry of microbial redox reactions*

Following the methodology of Rittman and McCarty 2001 (1), and subsequent modifications and applications for marine microbial ecosystem modeling (2, 3), we use $C_{c_{OM}}H_{h_{OM}}O_{o_{OM}}N_{n_{OM}}$ and $C_{c_B}H_{h_B}O_{o_B}N_{n_B}$ to describe compositions of OM and biomass (B), respectively. We define *d* to normalize redox reactions to one electron. To achieve mass balance (i.e., balance H while all the other atoms and electrons are balanced) in the equations:

$$d_{OM} = 4c_{OM} + h_{OM} - 2o_{OM} - 3n_{OM}$$
$$d_B = 4c_B + h_B - 2o_B - 3n_B$$

We assume a Redfieldian composition of OM ($C_{c_{OM}}H_{h_{OM}}O_{o_{OM}}N_{n_{OM}}$ = $C_{6.6}H_{10.9}O_{2.6}N$) (Anderson 1995), and all microbial functional types in the model have the same composition of biomass ($C_{c_B}H_{h_B}O_{o_B}N_{n_B}$ = $C_5H_7O_2N$) (Zimmerman et al. 2014). We define *f* as the fraction of electrons used in biomass synthesis. The biomass yield of each resource depends on *f*. For example, the yield of OM ($y_{OM}$) equals $\frac{d_{OM}}{d_B}f$ as shown in the equations below.

*Aerobic heterotrophs, $B_{het}$*

The half reactions are:

$$\frac{1}{d_{OM}}OM + \frac{2c_{OM} - o_{OM}}{d_{OM}}H_2O \rightarrow \frac{n_{OM}}{d_{OM}}NH_4^+ + \frac{c_{OM}}{d_{OM}}CO_2 + \frac{d_{OM} - n_{OM}}{d_{OM}}H^+ + e^-$$
$$(1-f) \times [\frac{1}{4}O_2 + H^+ + e^- \rightarrow \frac{1}{2}H_2O]$$
$$f \times [\frac{n_B}{d_B}NH_4^+ + \frac{c_B}{d_B}CO_2 + \frac{d_B - n_B}{d_B}H^+ + e^- \rightarrow \frac{1}{d_B}B_{N1234} + \frac{2c_B - o_B}{d_B}H_2O]$$

Ignoring $H_2O$ and $H^+$, the full reaction for $B_{het}$ is:

$$\frac{1}{d_{OM}}OM + \frac{1-f}{4}O_2 \rightarrow \frac{f}{d_B}B_{het} + \left(\frac{c_{OM}}{d_{OM}} - \frac{c_B f}{d_B}\right)CO_2 + \left(\frac{n_{OM}}{d_{OM}} - \frac{n_B f}{d_B}\right)NH_4^+$$

*Denitrifiers that reduce $NO_3^-$ to $N_2$, $B_{NO3-\rightarrow N2}$*

The half reactions are:

$$\frac{1}{d_{OM}}OM + \frac{2c_{OM} - o_{OM}}{d_{OM}}H_2O \rightarrow \frac{n_{OM}}{d_{OM}}NH_4^+ + \frac{c_{OM}}{d_{OM}}CO_2 + \frac{d_{OM} - n_{OM}}{d_{OM}}H^+ + e^-$$
$$(1-f) \times [\frac{1}{5}NO_3^- + \frac{6}{5}H^+ + e^- \rightarrow \frac{1}{10}N_2 + \frac{3}{5}H_2O]$$
$$f \times [\frac{n_B}{d_B}NH_4^+ + \frac{c_B}{d_B}CO_2 + \frac{d_B - n_B}{d_B}H^+ + e^- \rightarrow \frac{1}{d_B}B_{NO3-\rightarrow N2} + \frac{2c_B - o_B}{d_B}H_2O]$$

Ignoring $H_2O$ and $H^+$, the full reaction for $B_{NO3-\rightarrow N2}$ is:

$$\frac{1}{d_{OM}}OM + \frac{1-f}{5}NO_3^- \rightarrow \frac{f}{d_B}B_{NO3-\rightarrow N2} + \left(\frac{c_{OM}}{d_{OM}} - \frac{c_B f}{d_B}\right)CO_2 + \left(\frac{n_{OM}}{d_{OM}} - \frac{n_B f}{d_B}\right)NH_4^+ + \frac{1-f}{10}N_2$$

*Denitrifiers that reduce $NO_3^-$ to $N_2O$, $B_{NO3-\rightarrow N2O}$*

The half reactions are:

$$\frac{1}{d_{OM}}OM + \frac{2c_{OM} - o_{OM}}{d_{OM}}H_2O \rightarrow \frac{n_{OM}}{d_{OM}}NH_4^+ + \frac{c_{OM}}{d_{OM}}CO_2 + \frac{d_{OM} - n_{OM}}{d_{OM}}H^+ + e^-$$
$$(1-f) \times [\frac{1}{4}NO_3^- + \frac{5}{4}H^+ + e^- \rightarrow \frac{1}{8}N_2O + \frac{5}{8}H_2O]$$
$$f \times [\frac{n_B}{d_B}NH_4^+ + \frac{c_B}{d_B}CO_2 + \frac{d_B - n_B}{d_B}H^+ + e^- \rightarrow \frac{1}{d_B}B_{NO3-\rightarrow N2O} + \frac{2c_B - o_B}{d_B}H_2O]$$

Ignoring $H_2O$ and $H^+$, the full reaction for $B_{NO3-\rightarrow N2O}$ is:

$$\frac{1}{d_{OM}}OM + \frac{1-f}{4}NO_3^- \rightarrow \frac{f}{d_B}B_{NO3-\rightarrow N2O} + \left(\frac{c_{OM}}{d_{OM}} - \frac{c_B f}{d_B}\right)CO_2 + \left(\frac{n_{OM}}{d_{OM}} - \frac{n_B f}{d_B}\right)NH_4^+ + \frac{1-f}{8}N_2O$$

*Denitrifiers that reduce $NO_3^-$ to $NO_2^-$, $B_{NO3-\rightarrow NO2-}$*

The half reactions are:

$$\frac{1}{d_{OM}}OM + \frac{2c_{OM} - o_{OM}}{d_{OM}}H_2O \rightarrow \frac{n_{OM}}{d_{OM}}NH_4^+ + \frac{c_{OM}}{d_{OM}}CO_2 + \frac{d_{OM} - n_{OM}}{d_{OM}}H^+ + e^-$$

$$(1-f) \times [\frac{1}{2}NO_3^- + H^+ + e^- \rightarrow \frac{1}{2}NO_2^- + \frac{1}{2}H_2O]$$

$$f \times \left[\frac{n_B}{d_B}NH_4^+ + \frac{c_B}{d_B}CO_2 + \frac{d_B - n_B}{d_B}H^+ + e^- \rightarrow \frac{1}{d_B}B_{NO3-\rightarrow NO2-} + \frac{2c_B - o_B}{d_B}H_2O\right]$$

Ignoring $H_2O$ and $H^+$, the full reaction for $B_{NO3-\rightarrow NO2-}$ is:

$$\frac{1}{d_{OM}}OM + \frac{1-f}{2}NO_3^- \rightarrow \frac{f}{d_B}B_{NO3-\rightarrow NO2-} + \left(\frac{c_{OM}}{d_{OM}} - \frac{c_B f}{d_B}\right)CO_2 + \left(\frac{n_{OM}}{d_{OM}} - \frac{n_B f}{d_B}\right)NH_4^+ + \frac{1-f}{2}NO_2^-$$

*Denitrifiers that reduce $NO_2^-$ to $N_2$, $B_{NO2-\rightarrow N2}$*

The half reactions are:

$$\frac{1}{d_{OM}}OM + \frac{2c_{OM} - o_{OM}}{d_{OM}}H_2O \rightarrow \frac{n_{OM}}{d_{OM}}NH_4^+ + \frac{c_{OM}}{d_{OM}}CO_2 + \frac{d_{OM} - n_{OM}}{d_{OM}}H^+ + e^-$$

$$(1-f) \times [\frac{1}{3}NO_2^- + \frac{4}{3}H^+ + e^- \rightarrow \frac{1}{6}N_2 + \frac{2}{3}H_2O]$$

$$f \times [\frac{n_B}{d_B}NH_4^+ + \frac{c_B}{d_B}CO_2 + \frac{d_B - n_B}{d_B}H^+ + e^- \rightarrow \frac{1}{d_B}B_{NO2-\rightarrow N2} + \frac{2c_B - o_B}{d_B}H_2O]$$

Ignoring $H_2O$ and $H^+$, the full reaction for $B_{NO2-\rightarrow N2}$ is:

$$\frac{1}{d_{OM}}OM + \frac{1-f}{3}NO_2^- \rightarrow \frac{f}{d_B}B_{NO2-\rightarrow N2} + \left(\frac{c_{OM}}{d_{OM}} - \frac{c_B f}{d_B}\right)CO_2 + \left(\frac{n_{OM}}{d_{OM}} - \frac{n_B f}{d_B}\right)NH_4^+ + \frac{1-f}{6}N_2$$

*Denitrifiers that reduce $NO_2^-$ to $N_2O$, $B_{NO2-\rightarrow N2O}$*

The half reactions are:

$$\frac{1}{d_{OM}}OM + \frac{2c_{OM} - o_{OM}}{d_{OM}}H_2O \rightarrow \frac{n_{OM}}{d_{OM}}NH_4^+ + \frac{c_{OM}}{d_{OM}}CO_2 + \frac{d_{OM} - n_{OM}}{d_{OM}}H^+ + e^-$$

$$(1-f) \times [\frac{1}{2}NO_2^- + \frac{3}{2}H^+ + e^- \rightarrow \frac{1}{4}N_2O + \frac{3}{4}H_2O]$$

$$f \times [\frac{n_B}{d_B}NH_4^+ + \frac{c_B}{d_B}CO_2 + \frac{d_B - n_B}{d_B}H^+ + e^- \rightarrow \frac{1}{d_B}B_{NO2-\rightarrow N2O} + \frac{2c_B - o_B}{d_B}H_2O]$$

Ignoring $H_2O$ and $H^+$, the full reaction for $B_{NO2-\rightarrow N2O}$ is:

$$\frac{1}{d_{OM}}OM + \frac{1-f}{2}NO_2^- \rightarrow \frac{f}{d_B}B_{NO2-\rightarrow N2O} + \left(\frac{c_{OM}}{d_{OM}} - \frac{c_B f}{d_B}\right)CO_2 + \left(\frac{n_{OM}}{d_{OM}} - \frac{n_B f}{d_B}\right)NH_4^+ + \frac{1-f}{4}N_2O$$

*Denitrifiers that reduce $N_2O$ to $N_2$, $B_{N2O\rightarrow N2}$*

The half reactions are:

$$\frac{1}{d_{OM}}OM + \frac{2c_{OM} - o_{OM}}{d_{OM}}H_2O \rightarrow \frac{n_{OM}}{d_{OM}}NH_4^+ + \frac{c_{OM}}{d_{OM}}CO_2 + \frac{d_{OM} - n_{OM}}{d_{OM}}H^+ + e^-$$

$$(1-f) \times [\frac{1}{2}N_2O + H^+ + e^- \rightarrow \frac{1}{2}N_2 + \frac{1}{2}H_2O]$$

$$f \times [\frac{n_B}{d_B}NH_4^+ + \frac{c_B}{d_B}CO_2 + \frac{d_B - n_B}{d_B}H^+ + e^- \rightarrow \frac{1}{d_B}B_{N2O\rightarrow N2} + \frac{2c_B - o_B}{d_B}H_2O]$$

Ignoring $H_2O$ and $H^+$, the full reaction for $B_{N2O\rightarrow N2}$ is:

$$\frac{1}{d_{OM}}OM + \frac{1-f}{2}N_2O \rightarrow \frac{f}{d_B}B_{N2O\rightarrow N2} + \left(\frac{c_{OM}}{d_{OM}} - \frac{c_B f}{d_B}\right)CO_2 + \left(\frac{n_{OM}}{d_{OM}} - \frac{n_B f}{d_B}\right)NH_4^+ + \frac{1-f}{2}N_2$$

For chemoautotrophs, OM is not in their reactions, and their electron donors are inorganic nitrogen including ammonium and nitrite. We still assume the same biomass composition of $C_5H_7O_2N$.

*The non-adjacent denitrifier that reduce $NO_3^-$ to $NO_2^-$ or reduce $N_2O$ to $N_2$*

All denitrifying functional types survive in the model except for the functional type with non-adjacent steps. We describe this functional type following the method used to describe a facultatively anaerobic population (3). Briefly, this denitrifying functional type uses one of the two Ns (i.e., $NO_3^-$ or $N_2O$) that results in a larger growth rate under each condition. Following our approach, we also apply a penalty for this two-step pathway. As a result, this denitrifying functional type is outcompeted by its single-step counterpart and is competitively excluded in all model simulations. Thus, our analysis considers only the six denitrifying functional types that do have a niche space in the model (Fig. 1a). We speculate that this non-adjacent denitrifier type might be favored under more complex conditions such as fluctuations of its two N sources. Future experiments and modeling work will be needed to determine its regulation on the two sets of metabolisms.

*Ammonia-oxidizing organisms, $B_{AOO}$*

As in Zakem et al 2022 (4), the half reactions are:

$$\frac{1}{6}NH_4^+ + \frac{1}{3}H_2O \rightarrow \frac{1}{6}NO_2^- + \frac{4}{3}H^+ + e^-$$

$$(1-f) \times [\frac{1}{4}O_2 + H^+ + e^- \rightarrow \frac{1}{2}H_2O]$$

$$f \times [\frac{n}{d}NH_4^+ + \frac{c-n}{d}CO_2 + \frac{n}{d}HCO_3^- + H^+ + e^- \rightarrow \frac{1}{d}B_{AOO} + \frac{2c-o+n}{d}H_2O]$$

The full reaction for $B_{AOO}$ is:

$$\left(\frac{1}{6} + \frac{f}{d}\right)NH_4^+ + \frac{cf}{d}CO_2 + \frac{1-f}{4}O_2 \rightarrow \frac{f}{d}B_{AOO} + \frac{1}{6}NO_2^-$$

*Nitrite-oxidizing bacteria, $B_{NOB}$*

As in Zakem et al 2022 (4), the half reactions are:

$$\frac{1}{2}NO_2^- + \frac{1}{2}H_2O \rightarrow \frac{1}{2}NO_3^- + H^+ + e^-$$

$$(1-f) \times [\frac{1}{4}O_2 + H^+ + e^- \rightarrow \frac{1}{2}H_2O]$$

$$f \times [\frac{n}{d}NH_4^+ + \frac{c-n}{d}CO_2 + \frac{n}{d}HCO_3^- + H^+ + e^- \rightarrow \frac{1}{d}B_{NOB} + \frac{2c-o+n}{d}H_2O]$$

The full reaction for $B_{NOB}$ is:

$$\frac{1}{2}NO_2^- + \frac{f}{d}NH_4^+ + \frac{cf}{d}CO_2 + \frac{1-f}{4}O_2 \rightarrow \frac{f}{d}B_{NOB} + \frac{1}{2}NO_3^-$$

*Anaerobic ammonia-oxidizing bacteria, $B_{AMX}$*

As in Zakem et al 2020 (3), we use *x* to indicate the electron flow partitioning in anammox bacteria, and we use the empirically estimated stoichiometry of the anammox process (5, 6) to infer *f* for anammox. The half reactions are:

$$x \times [\frac{1}{3}NH_4^+ \rightarrow \frac{1}{6}N_2 + \frac{3}{4}H^+ + e^-]$$

$$(1-x) \times [\frac{1}{8}NH_4^+ \rightarrow \frac{1}{8}NO_3^- + \frac{1}{2}H^+ + e^-]$$

$$(1-f) \times [\frac{1}{3}NO_2^- + \frac{4}{3}H^+ + e^- \rightarrow \frac{1}{6}N_2 + \frac{2}{3}H_2O]$$

$$f \times [\frac{n}{d}NH_4^+ + \frac{c-n}{d}CO_2 + \frac{n}{d}HCO_3^- + H^+ + e^- \rightarrow \frac{1}{d}C_cH_hO_oN_n + \frac{2c-o+n}{d}H_2O]$$

The full reaction for anammox ($B_{AMX}$) is:

$$\left(\frac{x}{3} + \frac{1-x}{8} + \frac{f}{d}\right)NH_4^+ + \frac{1-f}{3}NO_2^- + \frac{cf}{d}CO_2 \rightarrow \frac{f}{d}B_{amx} + \frac{1-x}{8}NO_3^- + \frac{x+1-f}{6}N_2$$

Empirically estimated stoichiometry of the anammox process is (5, 6):

$$NH_4^+ + 1.146NO_2^- + 0.071HCO_3^- + 0.057H^+ \rightarrow 0.986N_2 + 0.161NO_3^- + 2.002H_2O + 0.071CH_{1.74}O_{0.31}N_{0.20}$$

We normalize the equation to one mole of N-based biomass:

$$5NH_4^+ + 5.73NO_2^- + 0.355HCO_3^- + 0.285H^+ \rightarrow 4.93N_2 + 0.805NO_3^- + 10.01H_2O + 0.071C_5H_{8.7}O_{1.55}N$$

$$70.423NH_4^+ + 80.704NO_2^- + 5HCO_3^- + 4.014H^+ \rightarrow 69.437N_2 + 11.338NO_3^- + 140.986H_2O + C_5H_{8.7}O_{1.55}N$$

Ignoring $H_2O$ and $H^+$, and using $CO_2$ instead of $HCO_3^-$, the equation becomes:

$$70.423NH_4^+ + 80.704NO_2^- + 5CO_2 \rightarrow C_5H_{8.7}O_{1.55}N + 11.338NO_3^- + 69.437N_2$$

Solving for *x*:

$$NO_3^- \div N_2 = \left(\frac{1-x}{8}\right) \div \left(\frac{x+1-f}{6}\right) \approx \left(\frac{1-x}{8}\right) \div \left(\frac{x+1}{6}\right) = \frac{11.338}{69.437}$$

$$x \approx 0.64$$

With *d* = 20, *c* = 5, *x* = 0.64, the full reaction for $B_{AMX}$ becomes:

$$\left(0.258 + \frac{f}{20}\right)NH_4^+ + \frac{1-f}{3}NO_2^- + \frac{f}{4}CO_2 \rightarrow \frac{f}{20}B_{amx} + 0.045NO_3^- + \frac{1.64-f}{6}N_2$$

$$\left(\frac{5.16}{f} + 1\right)NH_4^+ + \frac{20 \times (1-f)}{3f}NO_2^- + 5CO_2 \rightarrow B_{amx} + \frac{0.9}{f}NO_3^- + \frac{20 \times (1.64-f)}{6f}N_2$$

When $f = 0.07$, the stoichiometry of the reaction is similar to that of Lotti et al. (2014) with a focus on reproducing their ammonia demand since ammonia is usually the limiting resource for anammox bacteria:

$$74.71NH_4^+ + 88.57NO_2^- + 5CO_2 \rightarrow B_{amx} + 12.86NO_3^- + 74.76N_2$$

**Analysis of uncertainties in pathway length penalty**

Since measurements or other estimates of *P* are limited, we analyze the effect of different P values on biomass yields $y_{OM}$ and $y_N$ (Fig. S8). There are four critical *P* values (0, $P_{C1}$, $P_{C2}$, and $P_{C3}$) that result in three scenarios (Fig. S8). Since biomass yield ($y_{OM}$) and resource concentration (*R**, subsistence concentration that is introduced below) of microbes cannot be negative, we calculate the largest *P* value ($P_{C3}$) that ensures all $y_{OM}$ and $R^*_{OM}$ values equal or greater than zero. The critical value $P_{C1}$ is when $y_{OM}$ (and $R^*_{OM}$) of the two $NO_3^-$-reducing denitrifiers ($NO_3^- \rightarrow NO_2^-$ and $NO_3^- \rightarrow NO_2^- \rightarrow N_2O$) are equal. The critical value $P_{C2}$ is when $y_N$ (and $R^*_N$) of $NO_3^- \rightarrow NO_2^- \rightarrow N_2O$ and $NO_3^- \rightarrow NO_2^- \rightarrow N_2O \rightarrow N_2$ denitrifiers are equal. When *P* is smaller than $P_{C1}$ and greater than 0, $NO_3^- \rightarrow NO_2^-$ denitrifiers lose their advantage because their $R^*_{OM}$ is no longer the smallest. This lowers the abundance of $NO_3^- \rightarrow NO_2^-$ denitrifier population under oligotrophic conditions and hinders the accumulation of $NO_2^-$. Under this scenario, biodiversity and features of OMZs cannot be achieved. When *P* is larger than $P_{C2}$ and smaller than $P_{C3}$, neither $R^*_{OM}$ nor $R^*_N$ of $NO_3^- \rightarrow N_2$ denitrifiers is the smallest among the $NO_3^-$-reducing types. Thus, this type is always outcompeted by other $NO_3^-$-reducing types. Under this scenario, biodiversity and biogeochemical features cannot be achieved either.

Therefore, the scenario used in our model is defined by critical values $P_{C1}$ and $P_{C2}$. When *P* is greater than $P_{C1}$ and smaller than $P_{C2}$, the $y_{OM}$ of the shorter denitrification steps is larger than that of longer denitrification steps using the same N, and the $y_N$ of the shorter denitrification steps is smaller. A growth advantage for the shortest denitrification pathways under N-replete conditions is analogous to the carbon overflow metabolism or fermentation under C-replete conditions (7–9). The intrinsic tradeoff between different metabolisms (i.e., OM vs. N) allows different denitrifying functional types to survive under different nutrient conditions. We use a value of *P* within this scenario (P = 0.16, but variations only affect the magnitude of the OM supply required to favor denitrifiers with longer steps) (Fig. S9).

**Using a metabolic model of Marinobacter to estimate $y_{OM}$ for different denitrification steps**

To aid in the interpretation of the results, we simulated Flux Balance Analysis (FBA) (10) to estimate metabolic budgets (i.e., relative uptakes, secretions, and biomass yields) using a metabolic model of Marinobacter D2M19. The model possesses the whole denitrification pathway. For the reconstruction of the metabolic model, we employed CarveMe (11), integrating the 'gap-fill' option to ensure strain viability in two distinct media previously validated experimentally: marine broth supplemented with lactate as the exclusive carbon source, and further supplemented with either oxygen or nitrate. Subsequently, three reactions were manually introduced into the model: a nitrogen transport and a nitrogen exchange reaction to facilitate nitrogen production, and a nitrite reductase to enable the conversion of nitrite into nitric oxide. The final model comprised 1032 genes, 2074 reactions, 1392 metabolites, and 202 exchanges.

We simulated FBA (10) to estimate metabolic budgets for aerobic respiration, $NO_3^- \rightarrow NO_2^-$, $NO_3^- \rightarrow N_2O$, and $NO_3^- \rightarrow N_2$ (full denitrification). In brief, FBA returns flux distributions across all reactions within the metabolic model, optimizing for growth while adhering to steady-state conditions and constraints imposed by the composition of the growth medium. We simulated FBA in 59 MBL media with different carbon sources (see Supplementary Table 1 for a list of the carbon sources used). Specifically, the simulated MBL media consisted of $Ca^{2+}$, $Cl^-$, $Co^{2+}$, $Cu^{2+}$, $Fe^{2+}$, $Fe^{3+}$, $H_2O$, $H^+$, $K^+$, $Mg^{2+}$, $Mn^{2+}$, molybdate, $Na^+$, ammonium, nitrate, $Ni^{2+}$, phosphate, sulfate, $Zn^{2+}$, riboflavin, biotin, thiamine, ascorbic acid, pantothenate, folate, nicotinate, 4-aminobenzoic acid, pyridoxine, lipoic acid, NAD, thiamin pyrophosphate, cyanocobalamin and a carbon source. To simulate aerobic growth, oxygen was also included in the medium.

In all media tested, FBA results showed aerobic growth when both oxygen and $NO_3^-$ were available (as expected), and $NO_3^-$ reduction was observed in anaerobic conditions only. To estimate the metabolic budget for $NO_3^- \rightarrow N_2O$ and $NO_3^- \rightarrow N_2$, we forced there to be no intermediate N excretion. We conducted FBA using the COBRA Toolbox v0.25.0 (10, 11) and optimizing for growth rate.

**Sensitivity tests for the growth rate of aerobic heterotrophs and denitrifiers**

We follow recently published model parameterizations derived from observations (4, 12) and test the traits that are known to be uncertain. In particular, heterotrophs are known to have two distinct lifestyles (oligotrophy and copiotrophy). Oligotrophs have lower maximum uptake rates for nutrients and thus a lower maximum growth rate but higher substrate affinities. Copiotrophs have higher maximum uptake rates but lower substrate affinities. Since the lower oxygen tolerance of $NO_2^-$-reducing denitrifiers than $NO_3^-$-reducing denitrifiers results from interactions between heterotrophs and autotrophs, different heterotrophic lifestyles might affect these interactions. We parameterize our original heterotrophic functional types based on the high abundance of SAR11 (oligotrophic heterotrophs) in OMZs (13–16). We also test the effect of including a set of copiotrophic heterotrophs in the model on the original results, which have the capability to grow much faster than NOB and other functional types when substrate is available, such as at the onset of an OM pulse in our model. Including these rapidly growing copiotrophs does not change our conclusions, in particular, our conclusions about the ability of NOB to inhibit nitrite reduction resulting in the lower oxygen sensitivities associated with nitrate-consuming denitrification modules.

**Supplementary Figures**

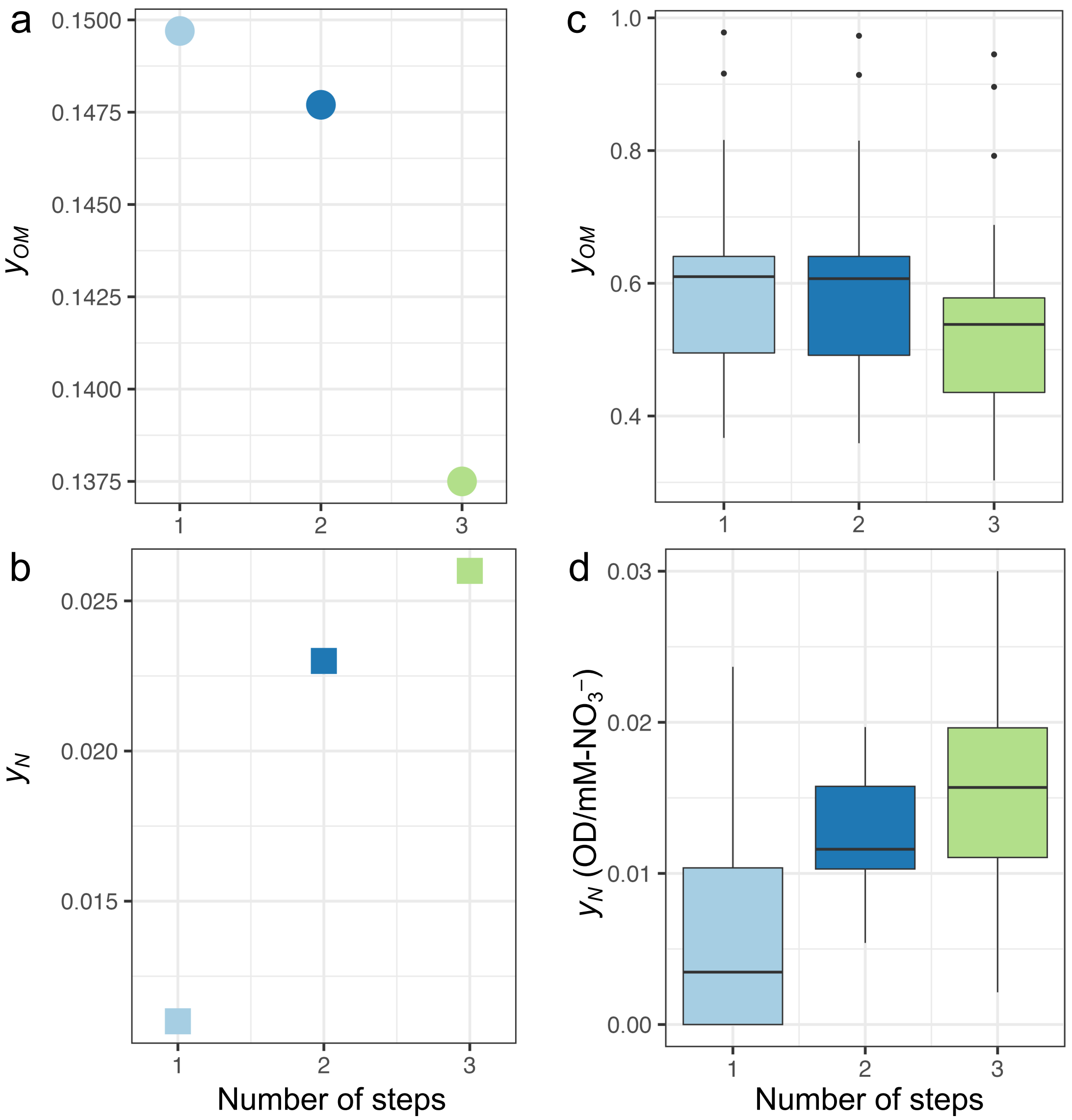


**Figure S1** The relationship between $y_{OM}$ and $y_N$ of $NO_3^-$-reducing denitrifiers and the number of denitrification steps (**a, b**) from our theoretical framework, (**c**) estimated with Flux Balance Analysis when simulating the growth of Marinobacter D2M19 in 59 different media, and (**d**) calculated from published data (17). The number of denitrification steps in **d** is determined following our theoretical framework: the presence of one or two genes of the same step is accounted as one step, and the NO reductase genes are ignored. We used data from 75 out of 78 isolates in the study containing genes encoding $NO_3^- \rightarrow NO_2^-$, $NO_3^- \rightarrow N_2O$, or $NO_3^- \rightarrow N_2$.

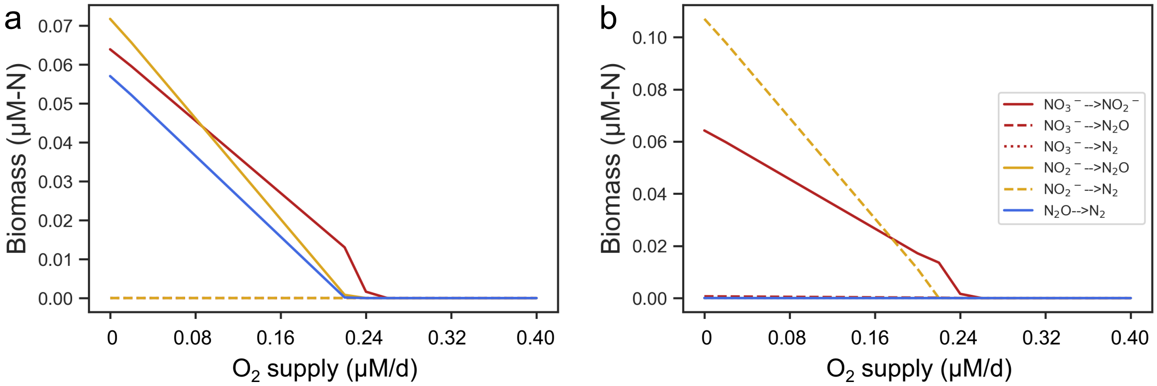


**Figure S2** Steady state solutions of the chemostat model with **(a)** only single-step denitrifiers ($NO_3^-$→$NO_2^-$, $NO_2^-$→$N_2O$, and $N_2O$→$N_2$) while setting the other denitrifiers' biomass to be zero and other microbes or **(b)** with six denitrifiers and other microbes. OM supply is 1 μM per day.

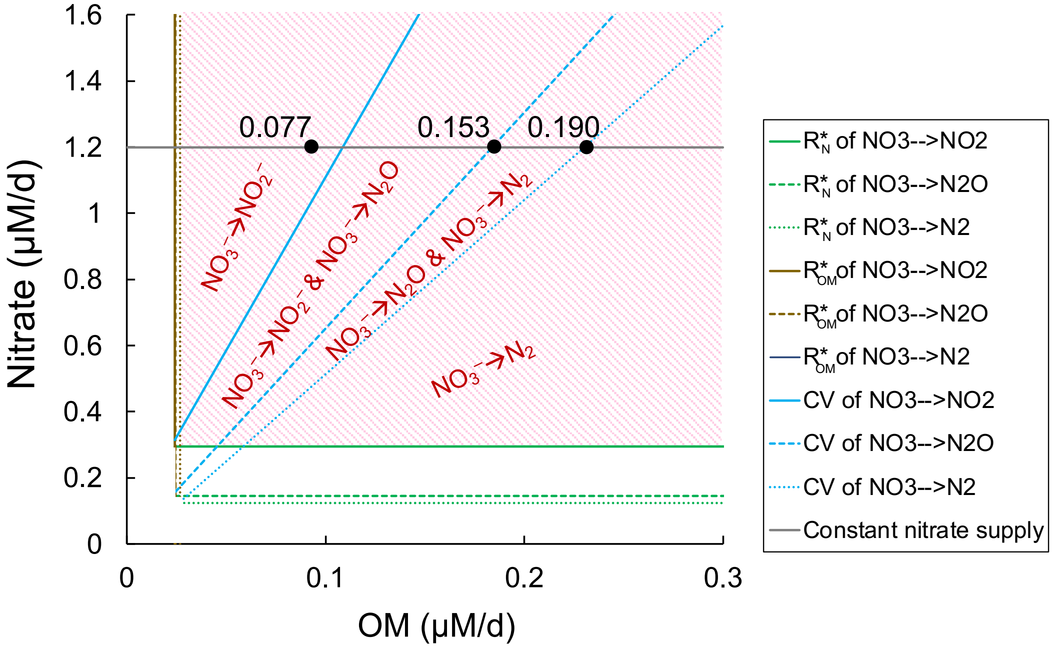


**Figure S3** Resource competition by three $NO_3^-$-reducing denitrifiers based on Tilman's resource-competition theory. The zero net growth isocline (ZNGI) of each species when $NO_3^-$ is limiting (i.e., $R^*_N$) is in green, and the ZNGI for OM limitation (i.e., $R^*_{OM}$) is in brown. Blue lines are consumption vectors (CV) of different denitrifiers. The pink dashed area highlights the space in which both nutrients ($NO_3^-$ and OM) exceed the subsistence concentration of resources (R*) of all species. Denitrifiers that survive in a certain resource space are indicated in red. The grey line is the constant $NO_3^-$ supply rate, and its intersections with CVs showing in black dots and labels are critical OM:$NO_3^-$ ratios, which define different survival regimes of microbes.

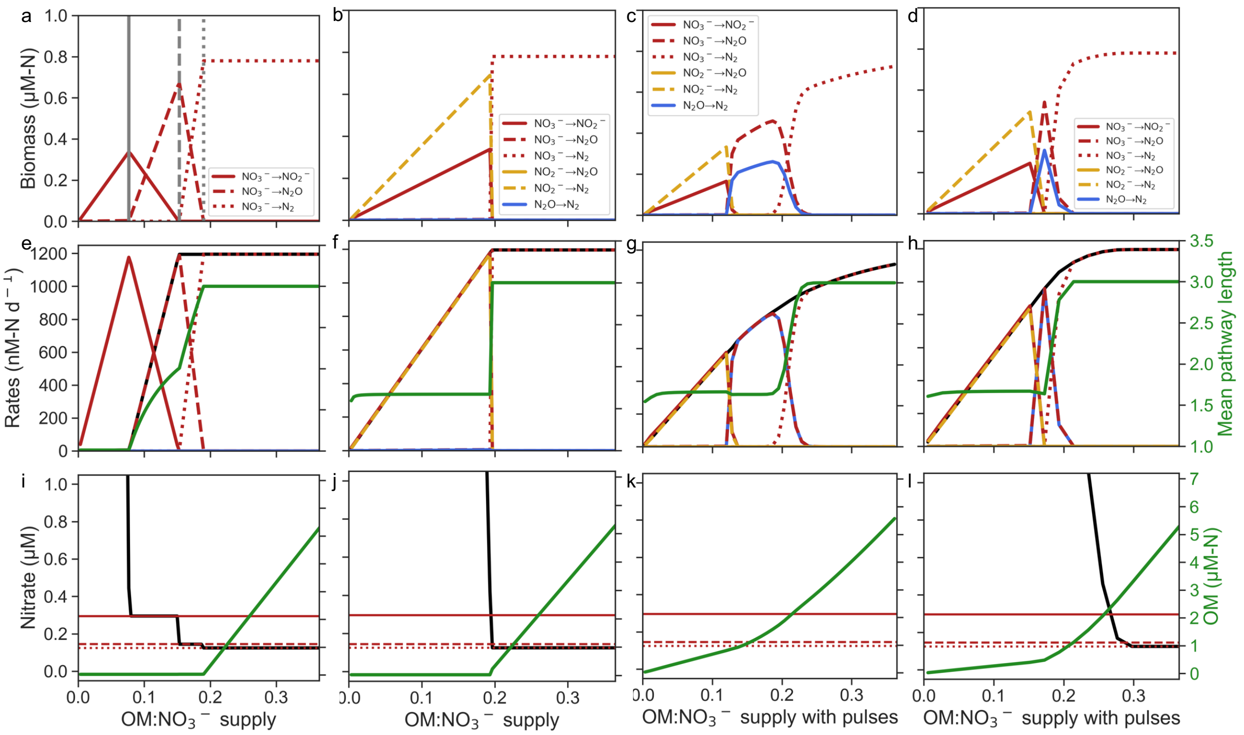


**Figure S4** The shift of denitrifier community composition (**a, b, c, d**), mean denitrification pathway length, N loss via denitrification and rate of each denitrifier type (**e, f, g, h**), and (quasi-)equilibrium $NO_3^-$ and OM concentrations and $R^*_N$ of each $NO_3^-$-reducing denitrifier type (**i, j, k, l**) with increasing OM:$NO_3^-$ supply when the $O_2$ supply rate is 0. Results from the model including (**a, e, i**) only three $NO_3^-$-reducing denitrifiers with constant OM supply, (**b, f, j**) only six denitrifier types with constant OM supply, six denitrifier types with OM pulses (**c, g, k**) every 50 days or (**d, h, l**) every 20 days. **a, b,** and **c** are replotted from **Fig. 2** for comparison with other plots.

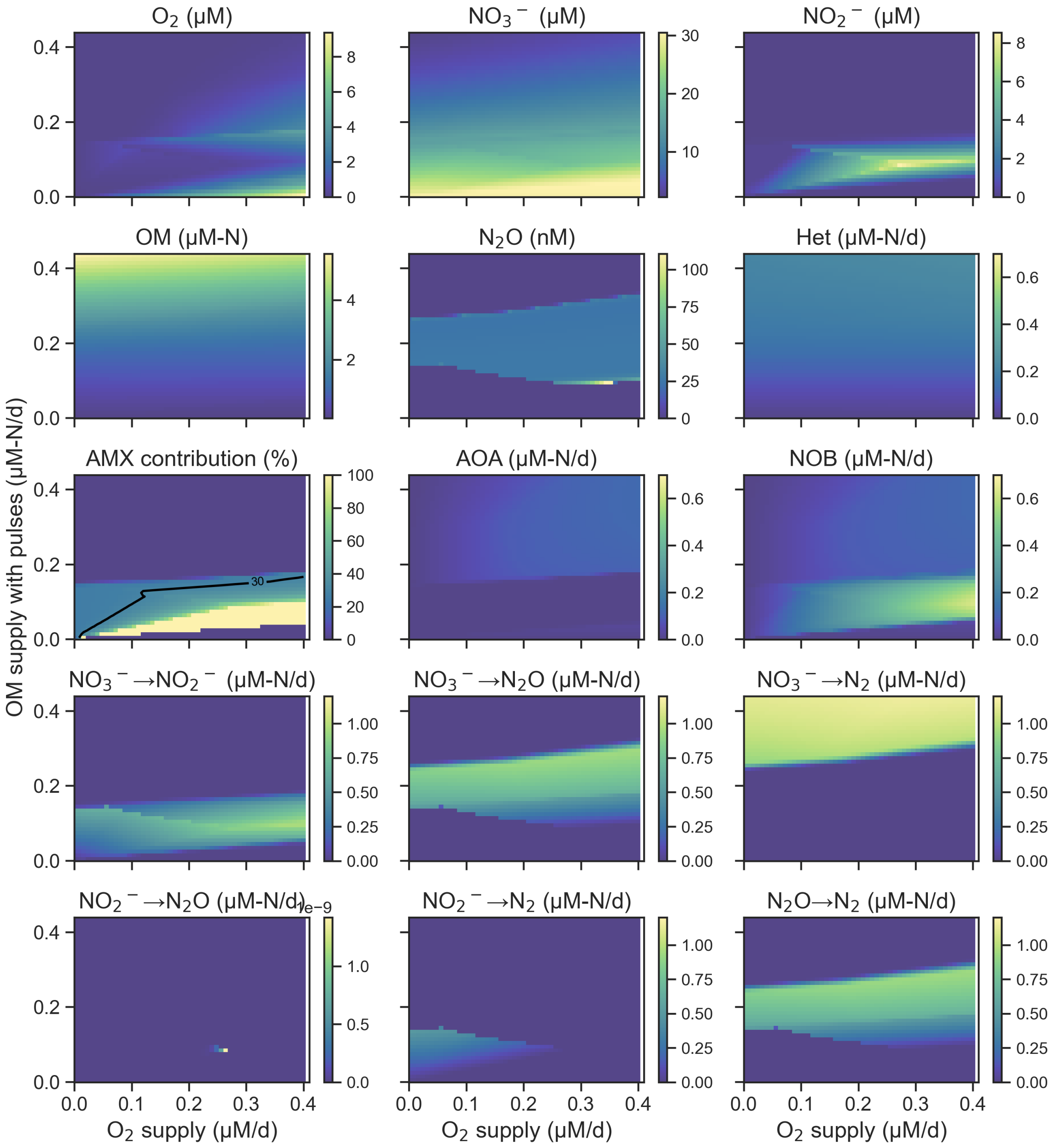


**Figure S5** Rates of all pathways and quasi-equilibrium concentrations of all nutrients estimated by the model with OM pulses. AMX (anammox) contribution (%) is the proportion of total $N_2$ produced from anammox, and the black contour line indicates the theoretical anammox contribution (30%) (18, 19).

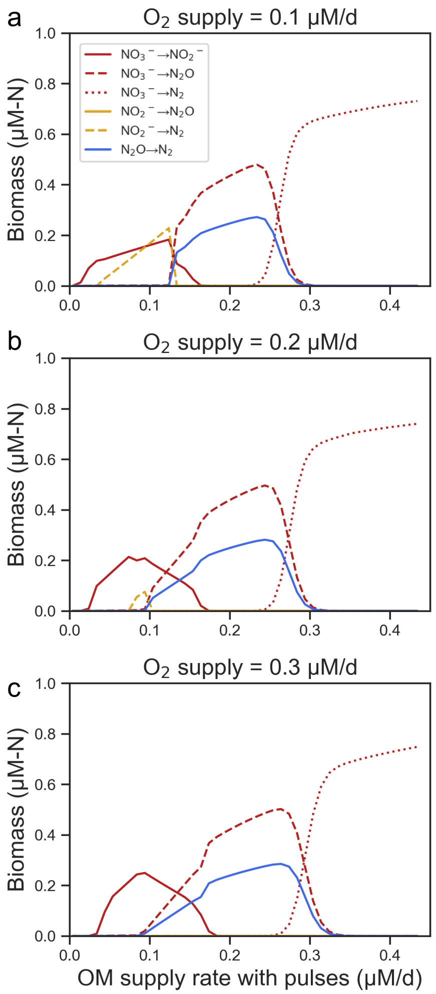


**Figure S6** The shift of denitrifier community composition with increasing OM supply rate when the $O_2$ supply rate is (**a**) 0.1, (**b**) 0.2, and (**c**) 0.3 μM/d. Results from the model including all six denitrifier types, anammox, AOA, NOB, and aerobic heterotrophs with OM pulses.

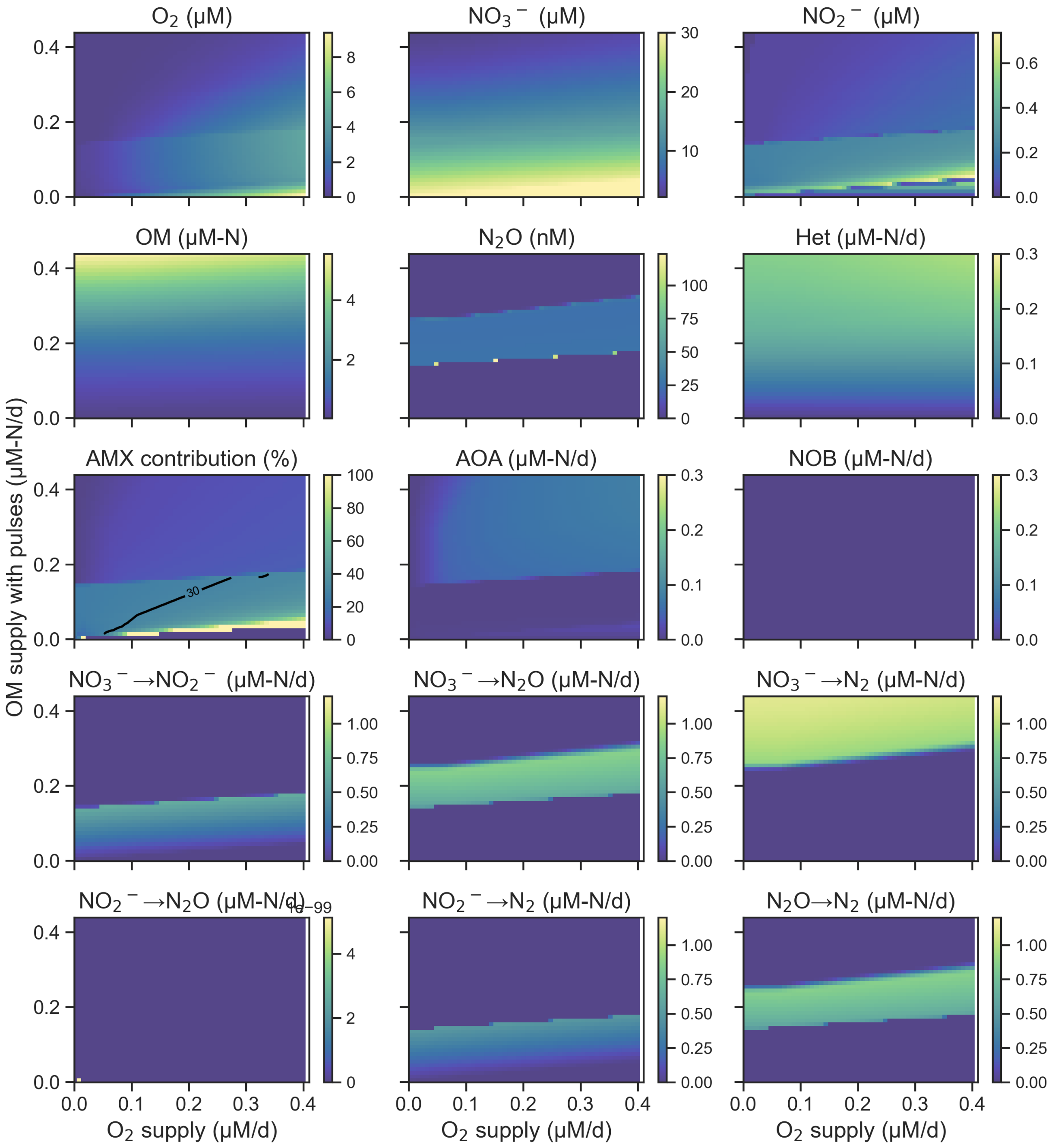


**Figure S7** Rates of all pathways and quasi-equilibrium concentrations of all nutrients estimated by the model with OM pulses when excluding NOB.

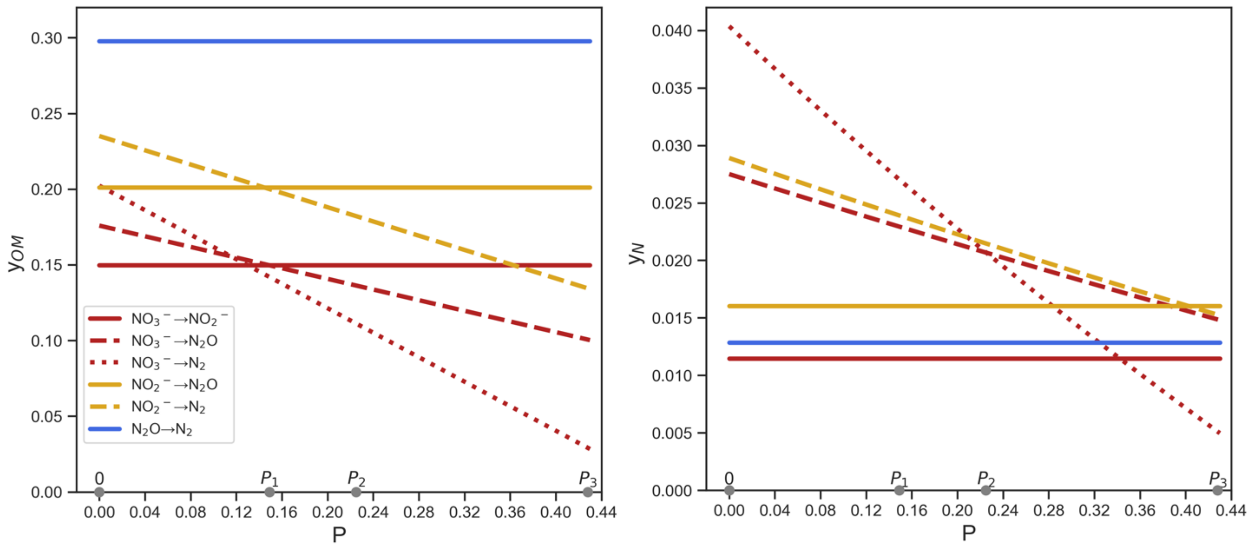


**Figure S8** The effect of step penalty on $y_{OM}$ and $y_N$ of denitrifiers. The same color indicates denitrifiers using the same N substrate, and the same line type indicates the same number of denitrification steps. Critical *P* values (0, $P_{C1}$, $P_{C2,}$ and $P_{C3}$) resulting in different scenarios are labeled on the x axis of each plot.

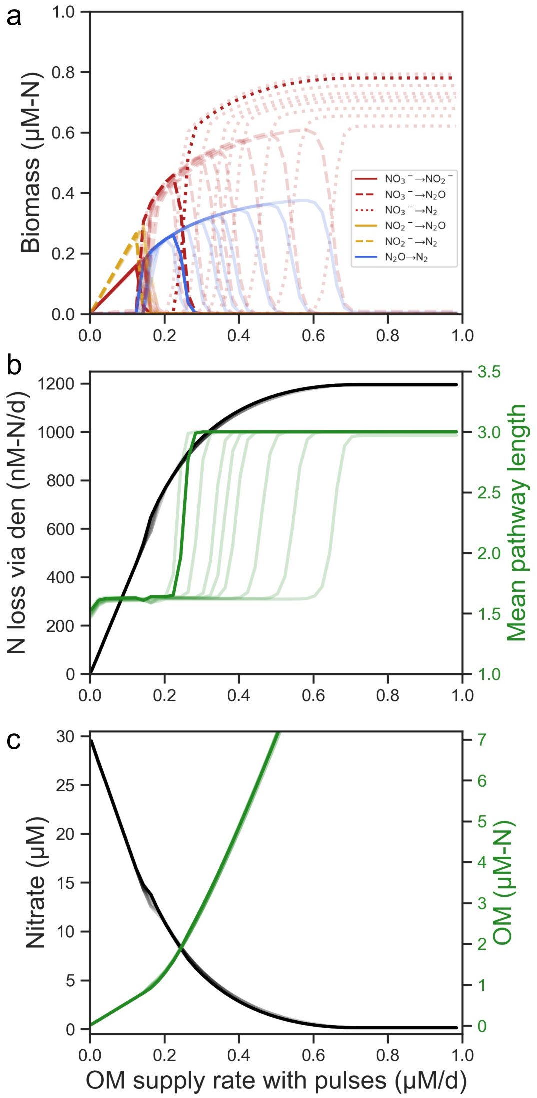


**Figure S9** The effect of *P* values on results from the model including all six denitrifier types, anammox, AOA, NOB, and aerobic heterotrophs with OM pulses. More transparent lines indicate *P* values varying from 0.155 to 0.220. The shift of denitrifier community composition (**a**), N loss via denitrification and mean denitrification pathway length (**b**), and quasi-equilibrium $NO_3^-$ and OM concentrations (**c**) with increasing OM supply rate when the $O_2$ supply rate is 0.

**Table S1** Results of metabolic modeling

| Metab id | Metab name | Carbon yield NO3->NO2 | Carbon yield NO3->NO2->N20 | Carbon yield NO3->NO2->N20->N2 |
|---|---|---|---|---|
| 12ppd__S_ | (S)-Propane-1,2-diol | 0.677 | 0.677 | 0.589 |
| glc__D_e | D-Glucose | 0.63 | 0.627 | 0.578 |
| 3amp_e | 3 AMP C10H12N5O7P | 0.816 | 0.815 | 0.792 |
| ala__L_e | L-Alanine | 0.542 | 0.539 | 0.479 |
| 4abut_e | 4-Aminobutanoate | 0.635 | 0.632 | 0.567 |
| acald_e | Acetaldehyde | 0.621 | 0.621 | 0.542 |
| ac_e | Acetate | 0.382 | 0.382 | 0.303 |
| mal__L_e | L-Malate | 0.419 | 0.419 | 0.375 |
| alaala_e | D-Alanyl-D-alanine | 0.559 | 0.556 | 0.498 |
| cys__L_e | L-Cysteine | 0.496 | 0.492 | 0.425 |
| gln__L_e | L-Glutamine | 0.492 | 0.489 | 0.435 |
| gly_e | Glycine | 0.367 | 0.359 | 0.309 |
| ser__L_e | L-Serine | 0.44 | 0.433 | 0.382 |
| thr__L_e | L-Threonine | 0.527 | 0.525 | 0.461 |
| arg__L_e | L-Arginine | 0.521 | 0.516 | 0.464 |
| cgly_e | Cys Gly C5H10N2O3S | 0.494 | 0.491 | 0.436 |
| malt_e | Maltose C12H22O11 | 0.613 | 0.61 | 0.558 |
| malttr_e | Maltotriose C18H32O16 | 0.618 | 0.616 | 0.565 |
| orn_e | Ornithine | 0.562 | 0.555 | 0.483 |
| asp__L_e | L-Aspartate | 0.459 | 0.452 | 0.414 |
| pro__L_e | L-Proline | 0.591 | 0.585 | 0.517 |
| meoh_e | Methanol | 0.569 | 0.56 | 0.485 |
| cit_e | Citrate | 0.415 | 0.411 | 0.37 |
| succ_e | Succinate | 0.445 | 0.444 | 0.386 |
| glu__L_e | L-Glutamate | 0.518 | 0.516 | 0.466 |
| dca_e | Decanoate (n-C10:0) | 0.592 | 0.592 | 0.49 |
| ddca_e | Dodecanoate (n-C12:0) | 0.637 | 0.637 | 0.538 |
| dha_e | Dihydroxyacetone | 0.61 | 0.607 | 0.557 |
| lac__D_e | D-Lactate | 0.542 | 0.539 | 0.479 |
| hdca_e | Hexadecanoate (n-C16:0) | 0.645 | 0.645 | 0.545 |
| hdcea_e | Hexadecenoate (n-C16:1) | 0.639 | 0.639 | 0.542 |
| ocdca_e | Octadecanoate (n-C18:0) | 0.648 | 0.648 | 0.547 |
| ocdcea_e | Octadecenoate (n-C18:1) | 0.642 | 0.642 | 0.544 |
| ttdca_e | Tetradecanoate (n-C14:0) | 0.642 | 0.642 | 0.542 |
| etoh_e | Ethanol | 0.752 | 0.752 | 0.672 |
| fald_e | Formaldehyde | 0.379 | 0.373 | 0.324 |
| fe3dcit_e | Fe(III)dicitrate | 0.411 | 0.407 | 0.365 |
| fru_e | D-Fructose | 0.63 | 0.627 | 0.578 |
| fum_e | Fumarate | 0.419 | 0.419 | 0.375 |
| g3pc_e | Sn-Glycero-3-phosphocholine | 0.978 | 0.973 | 0.945 |
| g3pe_e | Sn-Glycero-3-phosphoethanolamine | 0.707 | 0.704 | 0.642 |
| g3pg_e | Glycerophosphoglycerol | 0.732 | 0.726 | 0.668 |
| g3pi_e | Sn-Glycero-3-phospho-1-inositol | 0.916 | 0.914 | 0.896 |
| g3ps_e | Glycerophosphoserine | 0.628 | 0.622 | 0.575 |
| gal_e | D-Galactose | 0.621 | 0.619 | 0.568 |
| gthrd_e | Reduced glutathione | 0.51 | 0.507 | 0.454 |
| glcr_e | D-Glucarate | 0.409 | 0.409 | 0.362 |
| glyc3p_e | Glycerol 3-phosphate | 0.749 | 0.743 | 0.688 |
| glyc2p_e | Glycerol 2-phosphate | 0.679 | 0.673 | 0.607 |
| glyc_e | Glycerol | 0.732 | 0.726 | 0.668 |
| gthox_e | Oxidized glutathione | 0.51 | 0.507 | 0.454 |
| his__L_e | L-Histidine | 0.468 | 0.465 | 0.416 |
| hxa_e | Hexanoate (n-C6:0) | 0.541 | 0.541 | 0.441 |
| mal__D_e | D-Malate | 0.406 | 0.404 | 0.359 |
| malthx_e | Maltohexaose | 0.636 | 0.633 | 0.585 |
| maltpt_e | Maltopentaose | 0.63 | 0.627 | 0.578 |
| maltttr_e | Maltotetraose | 0.621 | 0.619 | 0.568 |
| man_e | D-Mannose | 0.63 | 0.627 | 0.578 |
| mnl_e | D-Mannitol | 0.683 | 0.678 | 0.625 |
